# Coherent activity at three major lateral hypothalamic neural outputs controls the onset of motivated behavior responses

**DOI:** 10.1101/2021.04.28.441785

**Authors:** Ekaterina Martianova, Alicia Pageau, Nikola Pausic, Tommy Doucet Gentiletti, Danahe Leblanc, Christophe D. Proulx

## Abstract

The lateral hypothalamus (LH) plays an important role in motivated behavior. However, it is not known how LH neural outputs dynamically signal to major downstream targets to organize behavior. We used multi-fiber photometry to show that three major LH neural outputs projecting to the dorsal raphe nucleus (DRN), ventral tegmental area (VTA), and lateral habenula (LHb) exhibit significant coherent activity in mice engaging motivated responses, which decrease during immobility. Mice engaging active coping responses exhibit increased activity at LH axon terminals that precedes an increase in the activity of serotonin neurons and dopamine neurons, indicating that they may play a role in initiating active responses stemming from LH signal transmissions. The optogenetic activation of LH axon terminals in either the DRN, VTA, or LHb was sufficient to increase mobility but had different effects on passive avoidance and sucrose consumption, suggesting that LH outputs use complementary mechanisms to control behavioral responses. Our results support the notion that the three LH neural outputs play complementary roles in initiating motivated behaviors.

## Introduction

An animal that wishes to maximize its survival must select optimal behavioral responses in order to correctly adapt to its environment. Deciding to execute or refrain from a specific action depends largely on the actual state of the environment that needs to be accurately integrated and then translated into adaptive responses (***Tye, 2018***; ***Verharen et al., 2020***). In other words, an animal will engage active behavioral responses to obtain a reward, avoid a punishment, or escape from an aversive context. Alternatively, it can engage passive coping behaviors to avoid being seen by a predator or to avoid exhaustion.

The lateral hypothalamus (LH) is a heterogeneous brain region that has been associated with a variety of behaviors related to motivation, reward, stress, arousal, and feeding (reviewed in ***Bonnavion et al.*** (***2016***) and ***Fakhoury et al.*** (***2020***)). More recently, its role in reward processing has been explored, specifically in innate defensive behaviors and in signaling aversive stimuli and the cues that promote them (***González et al., 2016a***; ***Hassani et al., 2016***; ***Noritake and Nakamura, 2019***; ***Karnani et al., 2020***; ***Lecca et al., 2017***; ***Trusel et al., 2019***; ***de Jong et al., 2019***).

Recent studies investigating individual LH neural outputs have shown that LH neuronal out-puts to the lateral habenula (LHb) and the dopaminergic (DA) ventral tegmental area (VTA) play important roles in these processes. LH neurons projecting to the lateral habenula (LH→LHb) are activated by aversive stimuli and encode cue-predicting aversive stimuli to motivate escape and avoidance (***Lecca et al., 2017***; ***Trusel et al., 2019***; ***Stamatakis et al., 2016***). LH projections to the dopaminergic ventral tegmental area (VTA) also play important roles in motivating compulsive reward seeking (***Nieh et al., 2015***), controlling innate defensive behaviors, and providing information on aversive outcomes (***de Jong et al., 2019***).

The dorsal raphe nucleus (DRN) is another brain center that plays an important role in reward processing and motivated behaviors (***Nakamura, 2013***). Direct activation of serotoninergic (5-HT, 5-hydroxytryptamine) DRN neurons increases active coping behavior (***Nishitani et al., 2019***), that originates from the prefrontal cortex and the LHb transmission (***Warden et al., 2012***; ***Amat et al., 2001***; ***Dolzani et al., 2016***). The DRN receives profuse projections from the LH (***Weissbourd et al., 2014***; ***Pollak Dorocic et al., 2014***; ***Zhou et al., 2017***; ***Ogawa and Watabe-Uchida, 2018***). Interestingly, pharmacological modulation of LH activity has been shown to exert either positive or negative control of serotoninergic DRN neurons (***Celada et al., 2002***), suggesting that it is involved in controlling DRN activity. However, the function of LH projections to the DRN in live behaving animal is unknown.

The LH, LHb, VTA and DRN are interconnected brain nuclei (***Weissbourd et al., 2014***; ***Pollak Dorocic et al., 2014***; ***Zhou et al., 2017***; ***Ogawa and Watabe-Uchida, 2018***; ***Stamatakis et al., 2016***; ***Lazaridis et al., 2019***; ***Lecca et al., 2017***). However, it not known how these structures are dynamically engaged to control behavior. Here, we used multi-fiber photometry (***Martianova et al., 2019***) to simultaneously monitor neural activity at LH axon terminals in three major brain targets. Our results show that largely non-overlapping LH neuronal populations transmit coherent signals to the DRN, VTA, and LHb. Activity along these pathways is coincident with the onset of motivated behavior responses. We also show that activity at LH axon terminals precedes activity at serotoninergic neurons in the DRN (*DRN*^5*HT*^) and dopaminergic neurons in the VTA (VTA^*DA*^). While optogenetic stimulation of LH terminals in the DRN, VTA, and LHb increased active coping behavior, it had a different effect on sucrose consumption, suggesting that these pathways play complementary roles in controlling behavior responses. Taken together, our results show that the three major LH neural outputs play complementary roles in engaging motivated behaviors.

## Results

### The LH→DRN, LH→VTA, and LH→LHb pathways are activated by aversive airpuffs and inhibited during sucrose consumption

To simultaneously examine activity at the LH→DRN, LH→VTA, and LH→LHb pathways (***Figure 1***A), we injected an adeno-associated virus (AAV) encoding the calcium indicator GCaMP6s (AAV-CAG-GCaMP6s) in the LH of wild-type mice. Optical fibers were implanted over the DRN, VTA, and LHb, allowing us to monitor activity at LH axon terminals projecting to the DRN, VTA, and LHb, respectively (***Figure 1***B). Four weeks post-injection, GCaMP6s expression was detected in the cell bodies of the LH and in the axon terminals of these fibers in the DRN, VTA, and LHb (***Figure 1***B). Calcium-dependent changes in fluorescence at axon terminals, which is a proxy for neural activity, was recorded simultaneously in all three pathways using multi-fiber photometry in freely behaving mice (***Figure 1***C) (***Martianova et al., 2019***). To directly compare the neural dynamics of these three LH neural outputs, the mice were placed in an open-field arena, and 1-s airpuffs (aversive stimulus) were delivered on top of the animals every 60 s (***Figure 1***D). ***Figure 1***E shows representative traces where spontaneous activity was detected at all three pathways, which significantly increased coincidentally with the delivery of the airpuffs (***Figure 1***F). Control mice injected with an AAV-eYFP exhibited no significant changes in fluorescence following the delivery of airpuffs (***Figure 1–Figure Supplement 1***C) indicating that changes in fluorescence are dependent on changes in calcium and GCaMP6s signals and do not result from movement artifacts.

**Figure 1.**
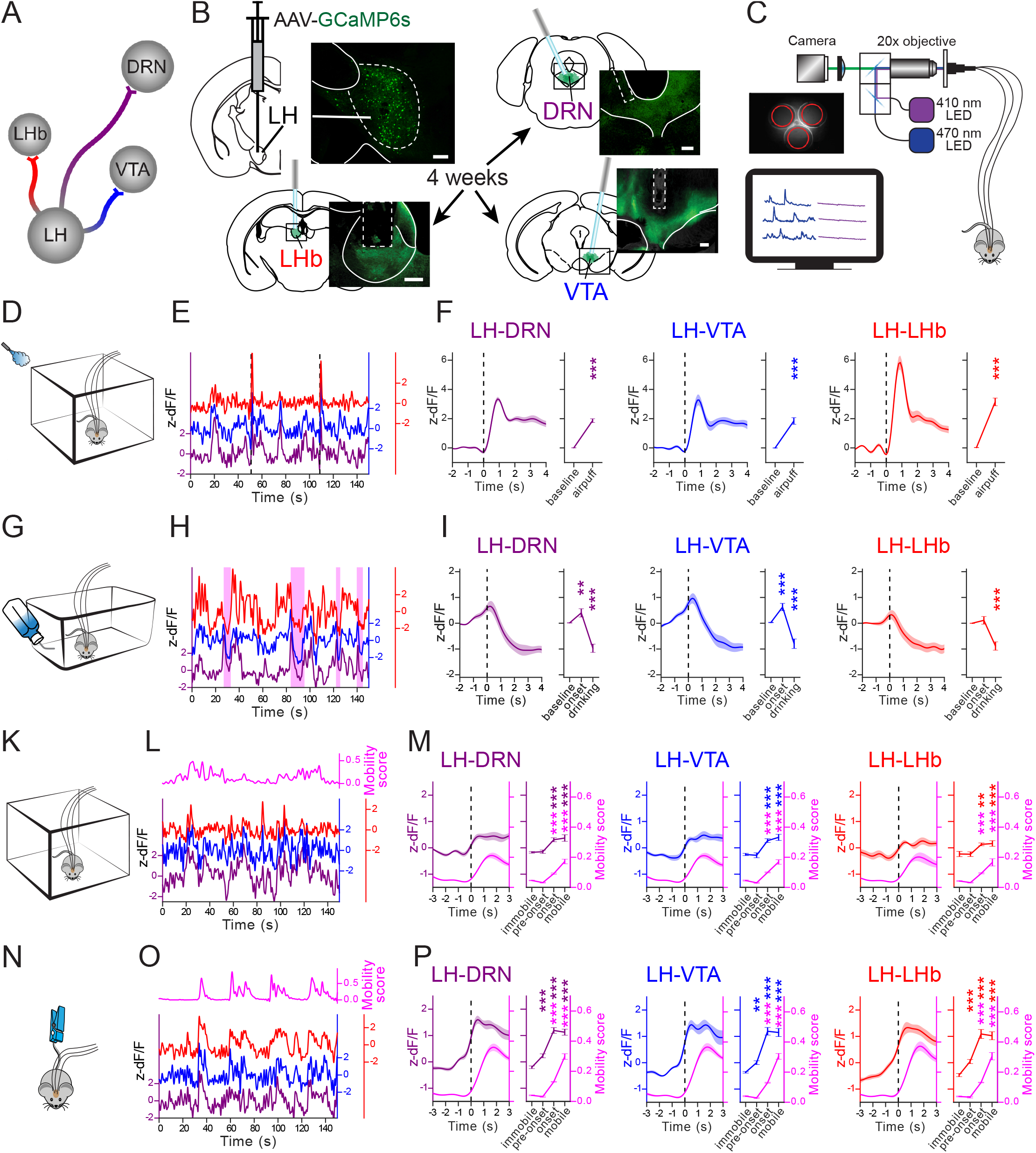
The LH→DRN, LH→VTA, and LH→LHb pathways are activated by airpuffs, at onset of mobility and are inhibited during sucrose consumption. (**A**) Diagram of the LH neural outputs targeted in this work. (**B**) Diagram of the experimental setup and fluorescence images of GCaMP6s expression in LH neurons and axon terminals at the DRN, VTA, and LHb (scale bar 200 *μ*m). (**C**) Diagram of the multi-fiber photometry calcium (*Ca*^2+^) imaging setup. (**D**) Experimental setup for the airpuffs. (**E**) Representative *Ca*^2+^ signal traces associated with the airpuffs (dashed vertical bars) simultaneously measured at the LH→DRN, LH→VTA, and LH→LHb pathways. (**F**) Peri-event plot of the average *Ca*^2+^ signal traces with all airpuff events at the LH→DRN, LH→VTA, and LH→LHb axon terminals. Plot of area under the curve (AUC) before and after the airpuffs. The lines represent means ± SEM (standard error of mean). Same convention as with **D-F** for the sucrose consumption test (**G-I**), the open field test (**K-M**), and the tail suspension test (**N-P**). Sucrose consumption events are represented by pink shaded box in **H**. The magenta lines are mobility scores. Repeated measures three-way ANOVA between factors group (GCaMP6s- and eYFP-expressing mice) and within factors pathway (LH→DRN, LH→VTA, and LH→LHb), and time period (different for each test) with post hoc Dunnett’s test. The *p* values were adjusted using the Bonferroni multiple testing correction method. \**p* < 0.05, \*\**p* < 0.01, \*\*\**p* < 0.001. **Figure 1–Figure supplement 1.** Recordings from the LH→DRN, LH→VTA, and LH→LHb pathways in control eYFP-expressing mice. **Figure 1–Figure supplement 2.** Pearson correlation between the signals recorded at the LH→DRN, LH→VTA, and LH→LHb pathways in mice expressing GCaMP6s and eYFP. **Figure 1–Figure supplement 3.** Cannulae placement in the mice expressing GCaMP6s **Figure 1–Figure supplement 4.** Cannulae placement in the mice expressing eYFP **Figure 1–Figure supplement 5.** The LH neural populations projecting to the DRN, VTA, and LHb are largely distinct populations. **Figure 1–Figure supplement 6.** *Ca*^2+^ imaging from the LH neurons projecting to the DRN and the VTA **Figure 1–Figure supplement 7.** The cannulae placement in mice prepared using the intersectional viral strategy **Figure 1–source data 1.** fig1_LH_apt-sct-oft-tst.xlsx contains AUC from APT, SCT, OFT, and TST experiments with mice expressing GCaMP6s and correlation analysis between outputs; related to figure 1 and figure supplement 2. **Figure 1–source data 2.** fig1-supp1_LH-eYFP.xlsx contains AUC from all experiments with control mice expressing eYFP and correlation analysis between outputs; related to figure supplement 1 and 2. **Figure 1–source data 3.** fig1-supp6_LHretro.xlsx contains AUC from all experiments with mice prepared using intersectional viral strategy; related to figure supplement 6. **Figure 1–source data 4.** Fig1_stats.ipynb contains statistical analysis related to figure 1 and its supplements.

To investigate how these pathways are modulated in an appetitive context, the same mice were water-deprived for 24 h and were then given free access to a 2% sucrose solution for 10 min (***Figure 1***G). In these sessions of sucrose consumption (SCT), the mice readily learned to drink from the sucrose dispenser. Drinking events were automatically detected with a lickometer, and are represented by pink boxes in the representative traces shown in ***Figure 1***H). Aligning the signals with the onset of drinking events revealed that activity at the LH→DRN and LH→VTA pathways increased significantly immediately prior to the onset of sucrose consumption, indicating that these pathways play a role in initiating motivated goal-directed behavior. This change was not seen with the LH→LHb pathway. In all three pathways, however, activity significantly decreased during sucrose consumption (***Figure 1***G). No specific changes in the signal at the onset of or during sucrose consumption were observed in eYFP-expressing control mice (***Figure 1–Figure Supplement 1***F).

### A Coherent increase in activity at the LH→DRN, LH→VTA, and LH→LHb pathways at the onset of mobility

Previous studies have shown that the LH→VTA and LH→LHb pathways play a role in motivating defensive behaviors. However, our results suggest that the LH neural outputs may also play a role in controlling spontaneously motivated behavior. For example, the mice fled when given an aversive airpuff, increased their mobility to reach a drinking spout, and stayed put during consumption.

To directly examine the role of LH neural outputs in motivated behavior, activity was measured at LH neural outputs and movements were tracked with an automated ANY-maze video tracking system. The mice were free to explore either an open field (OFT) (***Figure 2***A-C) or were suspended by the tail (tail suspension test, TST) (***Figure 2***D-F). OFT and TST are neutral and stressful contexts, respectively, and presumably engage different motivational drives to move. ***Figure 1***L and O show representative activity traces monitored at the LH neural outputs and superimposed on the mobility score (magenta line), of a mouse placed in the open field or a mouse suspended by its tail, respectively. In these contexts, activity at the LH→DRN, LH→VTA, and LH→LHb pathways significantly increased at mobility onset. In addition, mice placed in the TST exhibited a significant increase in activity at all three LH neural outputs immediately prior to the onset of mobility. A Pearson correlation analysis confirmed the coherence between these pathways, which we did not observe in eYFP-expressing mice (***Figure 1–Figure Supplement 2***). Taken together, these results show that these three major LH neural outputs exhibit coherent activity.

**Figure 2.**
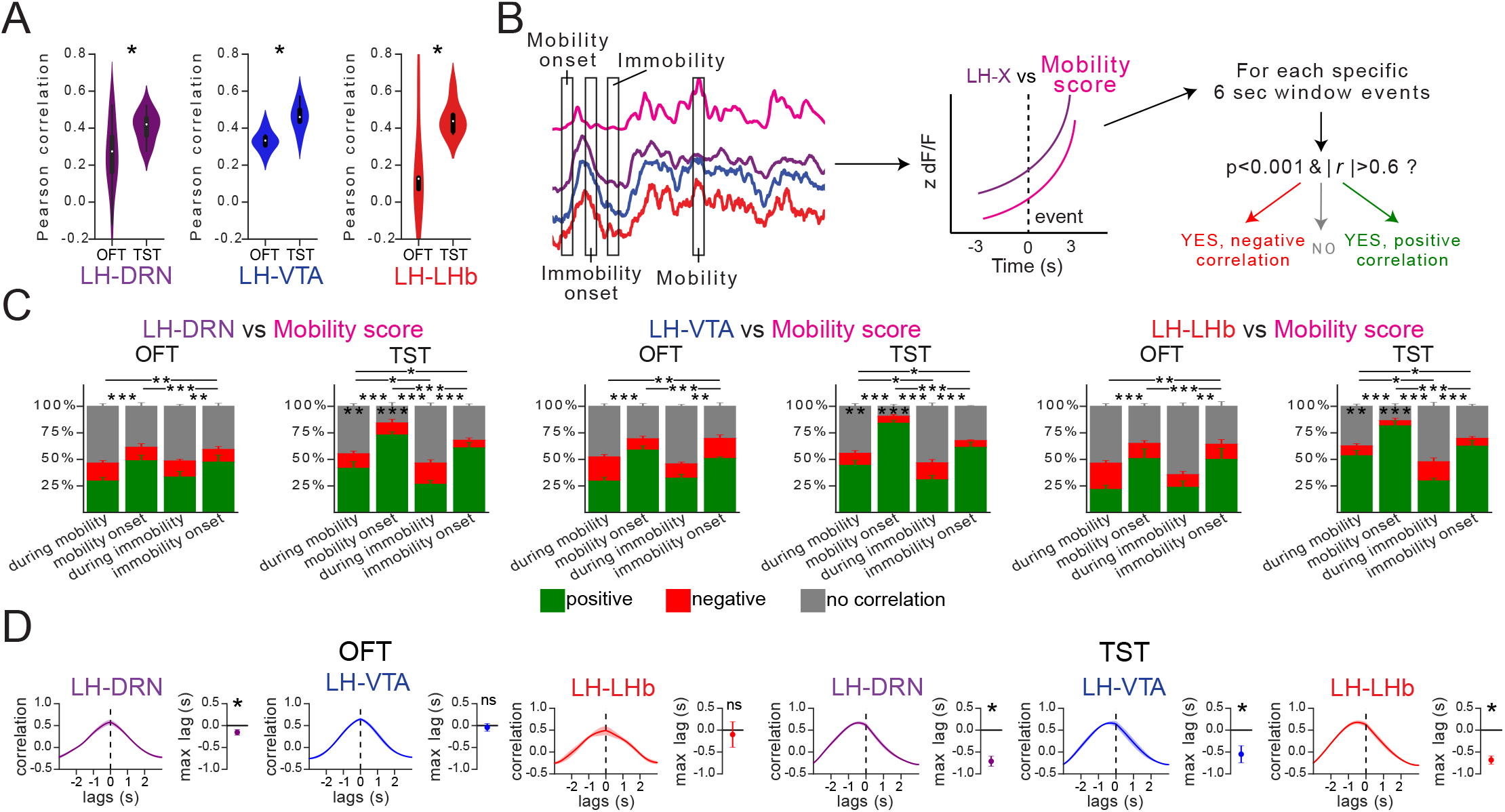
There was a high correlation between activity and mobility score at the LH→DRN, LH→VTA, and LH→LHb pathways during the TST. (**A**) Pearson correlation between *Ca*^2+^ signal at the LH→DRN, LH→VTA, and LH→LHb pathways, and mobility scores during the OFT or TST. One sample t-test. Three-way mixed ANOVA between factors group (GCaMP6s- and eYFP-expressing mice), and within factors test (OFT and TST) and pathways (LH→DRN, LH→VTA, and LH→LHb) with post hoc Tukey test. The *p* values were adjusted using the Bonferroni multiple testing correction method. (**B**) Schematic of the event selection. Events at the onset mobility and immobility, and random events during mobility and immobility were chosen, and the Pearson correlation at 6 seconds peri-events between the *Ca*^2+^ signal and the mobility score was calculated. Correlations with p < 0.001 and r > 0.6 were considered as positive, p < 0.001 and r < 0.6 as negative, and the others as uncorrelated. (**C**) Fraction of positive (green), negative (red), and uncorrelated events (gray) in the OFT and TST for the LH→DRN, LH→VTA, and LH→LHb pathways. Four-way MANOVA between the factors group (GCaMP6s- and eYFP-expressing mice), test (OFT and TST), and pathways (LH→DRN, LH→VTA, and LH→LHb), and within factor events (during mobility, mobility onset, during immobility, immobility onset) for positive and negative correlations with a post hoc four-way mixed ANOVA and Tukey test for multiple comparisons. The *p* values were adjusted using the Bonferroni multiple testing correction method. The main effect was due to differences in the number of positive correlations. There was no difference in factor output and its interactions with other factors. Asterisks on the bars show the difference between tests. (**D**) Cross-correlation analysis between the *Ca*^2+^ signal and the mobility score at mobility onset peri-events. The lines represent means ± SEM of correlations vs. lag times and means ± SEM of lag times with maximum correlation. One sample t-test. Two-way ANOVA within factors tests (OFT and TST) and pathways (LH→DRN, LH→VTA, and LH→LHb) with post hoc Tukey test. The *p* values were adjusted using the Bonferroni multiple testing correction method. \**p* < 0.05, \*\**p* < 0.01, \*\*\**p* < 0.001. **Figure 2–Figure supplement 1.** Correlation analysis between the signal at one of the LH neural output pathways and mobility score in mice expressing eYFP. **Figure 2–source data 1.** fig2_LH_movementCorrelation.xlsx contains correlation results of GCaMP6s-expressing mice; related to figure 1. **Figure 2–source data 2.** fig2_LH-eYFP_movementCorrelation.xlsx contains correlation results of eYFP-expressing mice; related to figure 1. **Figure 2–source data 3.** Fig2_stats.ipynb contains statistical analysis related to figure 2 and its supplements.

### Distinct LH neuronal populations project to DRN, VTA, and LHb

The coherence observed at the LH neural outputs could be explained if there were extensive collateral projections from the LH to the DRN, VTA, and LHb. To determine whether LH neural outputs send out significant collateral projections, we first injected the same mouse with three fluorescently labeled retrograde markers in the DRN, VTA, and LHb. Three to four days later, we performed a histological analysis and quantified LH neurons that were positive for one or more fluorescent markers (***Figure 1–Figure Supplement 5***). Our analysis revealed largely non-overlapping LH neuronal populations projecting either to the DRN, the VTA, or the LHb, with less than 5% of LH neurons being positive for two or three fluorescent markers. This observation is consistent with a previous report showing that distinct LH neural outputs projecting to VTA, LHb and dPAG (***de Jong et al., 2019***). To further determine whether the activity measured at LH axon terminals was from distinct LH populations, we used an intersectional viral approach to express the red-shifted calcium sensor jrGECO1a (***Dana et al., 2016***), and GCaMP6s in DRN- and VTA-projecting LH neurons, respectively (***Figure 1–Figure Supplement 6***A) (***Martianova et al., 2019***; ***Kakava-Georgiadou et al., 2019***). To this end, AAV vectors with retrograde properties (***Tervo et al., 2016***), encoding the cre or FlpO recombinase were injected into the DRN and the VTA, respectively. A cre-dependent AAV encoding jrGECO1a (AAV-DIO-jrGECO1a) and a FlpO-dependent AAV encoding GCaMP6s (AAV-fDIO-GCaMP6s) were injected into the LH of the same mice. As with fluorescent retrograde markers, this approach mainly labelled non-overlapping, and distinct LH neuronal populations (***Figure 1–Figure Supplement 6***A). An optical fiber was implanted over the LH of these mice to monitor signals from jrGECO1a and GCaMP6s expressed in DRN- and VTA-projecting LH neurons by dual-color fiber photometry (***Martianova et al., 2019***). When the mice were tested using the same behaviors described above, the calcium signals perfectly replicated the results previously obtained from the axon terminals recordings. These results indicate that the LH sends out largely independent projections, that it signals to the DRN, VTA and LHb, and that this activity is coherent to mobility onset.

### Increased activity at the LH→DRN, LH→VTA, and LH→LHb pathways precedes coping responses

A Pearson correlation analysis between LH neural output activity and mobility score measured with the OFT and TST showed a significant positive correlation, with the exception of the LH→LHb pathway with the OFT (***Figure 2***A), indicating that these pathways play a role in controlling movement. This correlation was significantly higher with the TST, an aversive context engaging active and passive coping responses (***Commons et al., 2017***). To further investigate whether the high correlation between activity and the mobility score could be attributed to specific time events during the OFT and TST, we performed peri-event correlation analyses at specific time points (events). Events of four types were chosen: at mobility onset, at immobility onset, and at random time points during mobility and immobility. For each event, a Pearson correlation was measured for each 6-seconds peri-event traces of the mobility score and the calcium signal recorded at each of the LH outputs. Correlations with *p* < 0.001 and *r* > 0.6 were considered as positive correlation events, *p* < 0.001 and *r* < 0.6 as negative correlation events, and the others events as having no correlation. The number of positive, negative, and no correlations events were counted for each mouse, each time event, and each behavior test. The means ± SEM for all of the animals are given in ***Figure 2***C. The statistical analysis revealed that the main difference between the groups was the number of positive correlations. The results were independent of the output (no difference by this factor or interactions with others). In other words, the number of positive correlations events was similar for the LH→DRN, LH→VTA, and LH→LHb pathways. This analysis revealed that the number of positive events was significantly higher at mobility onset and during mobility in the TST than in the OFT. The highest number of positive correlations events was observed at mobility onset during the TST. A cross-correlation analysis between activity and mobility score also revealed that changes in activity from all three LH neural outputs preceded the change in mobility at mobility onset when the mice initiated an active coping response in the TST (***Figure 2***D). These results indicate that there is a significant correlation in activity between these three LH neural outputs, and suggest that activity at these LH neural outputs controls motivated behavior, particularly to engage a coping response in a stressful context.

### Increased activity at LH axon terminals precedes increased activity of serotonin and dopamine neurons

To determine whether the change in activity measured at LH axon terminals was correlated with a postsynaptic change in activity, we have performed dual-color fiber photometry recordings of LH axon terminals while co-monitoring the activity of serotonin and dopamine neurons in the DRN and VTA respectively. To this end, a cre-dependent AAV encoding jrGECO1a (AAV-DIO-jrGECO1a) was injected into the DRN of ePet-cre mice, or in the VTA of DAT-ires-cre mice. These are transgenic mouse lines exclusively expressing the recombinase cre in 5-HT neurons (***Scott et al., 2005***) and DA neurons (***Bäckman et al., 2006***), respectively. AAV-GCaMP6s was also injected into the LH of the same mice, and optical fibers were implanted in the DRN and the VTA as described above (***Figure 3***A, D). Four weeks post-injection, GCaMP6s expression was observed in the LH axons terminals and jrGECO1a expression was observed in the cell bodies of 5-HT neurons (Tph2+) and DA neurons (TH+) in the DRN and VTA, respectively (***Figure 3–Figure Supplement 1***A, D). This approach allowed us to simultaneously monitor activity in LH axons terminals (green fluorescence from GCaMP6s) and in genetically-defined 5-HT and DA neurons (red fluorescence from jrGECO1a) using the same optical fiber. Representative traces show spontaneous activity from LH axon terminals as well as from DRN^5*HT*^ and VTA^*DA*^ neurons. The activity at DRN^5*HT*^ neurons significantly increased with aversive airpuffs and decreased during sucrose consumption, replicating the results from the LH axon terminals (***Figure 3–Figure Supplement 1***B, E). However, the activity of VTA^*DA*^ neurons increased significantly with aversive airpuffs and during sucrose consumption (***Figure 3–Figure Supplement 1***C, F). When evaluated by the OFT and TST, the activity of VTA^*DA*^ neurons increased significantly at the onset of mobility in the TST, but not in the OFT (***Figure 3***B, C). However, the activity of DRN^5*HT*^ increased significantly at mobility onset in both tests (***Figure 3***E, F). In addition, the activation of DRN^5*HT*^ and VTA^*DA*^ neurons was preceded by the activation of the LH→DRN and the LH→VTA pathways (***Figure 3***C, F). These results confirm that axon terminal activity is correlates with the postsynaptic activity of DRN^5*HT*^ and VTA^*DA*^ neurons, suggesting that the LH may be an important relay for transmitting signals to serotoninergic and dopaminergic nuclei in order to engage motivated behaviours in aversive contexts.

**Figure 3.**
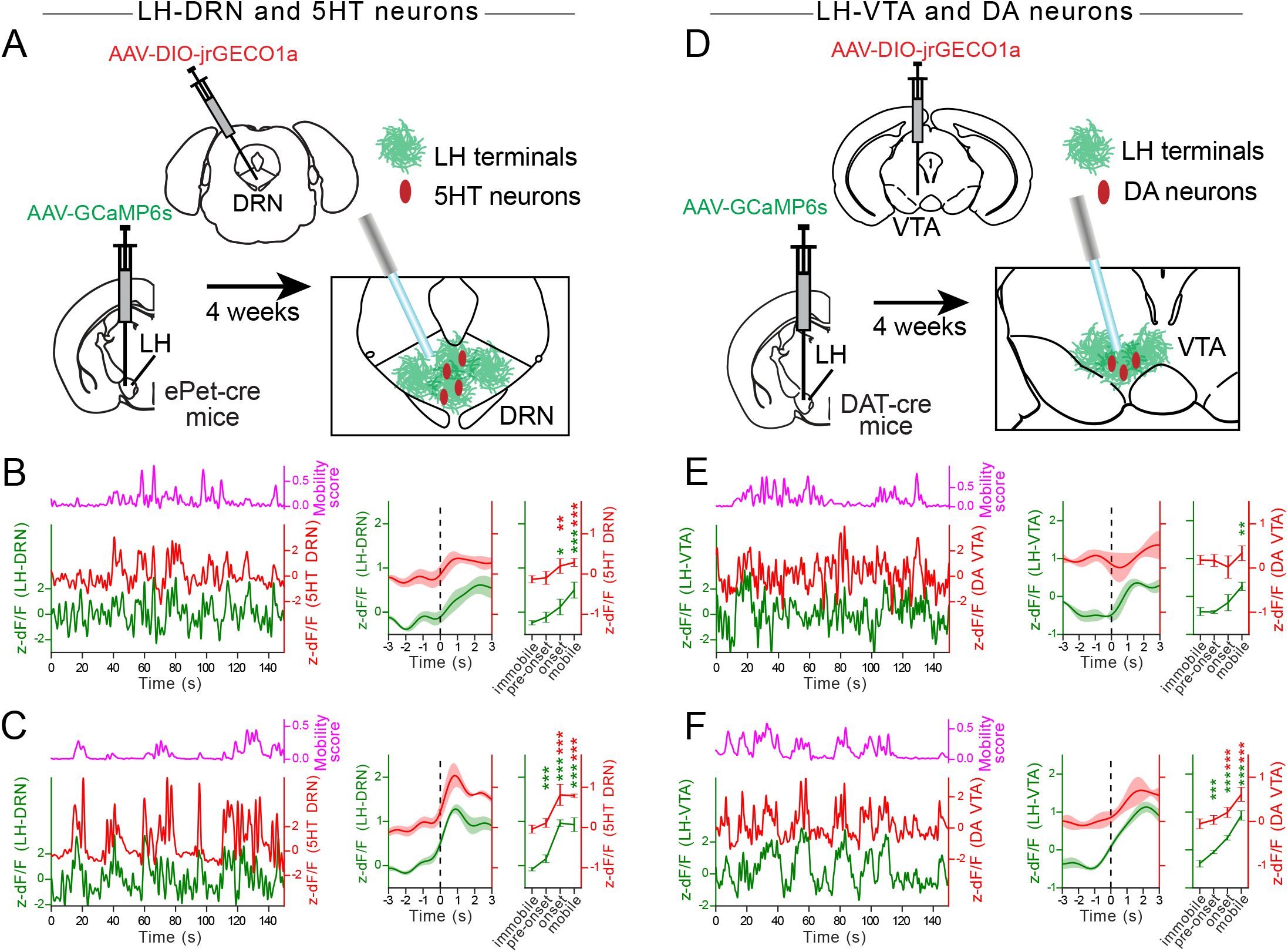
Increased activity of DRN^5*HT*^ and VTA^*DA*^ neurons follows increased activity in the LH→DRN and LH→VTA pathways at mobility onset in the TST. (**A, D**) Diagrams of the experimental setups for simultaneous recordings from the DRN^5*HT*^ neurons and the LH→DRN pathway (**A**) and from the VTA^*DA*^ neurons and the LH→VTA pathway (**D**). (**B, E**) Representative *Ca*^2+^ signal traces from the DRN^5*HT*^ neurons and the LH→DRN pathway(**B**) and the VTA^*DA*^ neurons and the LH→VTA pathway (**E**) during the OFT (**left**). Peri-event plots of the average *Ca*^2+^ signals at all mobility onsets and plots for AUC during immobility and mobility, and at mobility pre-onset and onset (**right**). The lines represent the means ± SEM (standard error of mean). Same convention as with **B, E** for the TST (**C, F**). the magenta lines are mobility scores. Repeated measures two-way ANOVA within factors pathways (DRN^5*HT*^ and LH→DRN, or VTA^*DA*^ and LH→VTA) and time periods (during immobility and mobility, at mobility pre-onset and onset) with post hoc Dunnett’s test. The *p* values were adjusted using the Bonferroni multiple testing correction method. \**p* < 0.05, \*\**p* < 0.01, \*\*\**p* < 0.001. **Figure 3–Figure supplement 1.** Simultaneous recordings at the DRN^5*HT*^ neurons and the LH→DRN pathway, and at the VTA^*DA*^ neurons and the LH→VTA pathway in the AP and the SCT. **Figure 3–Figure supplement 2.** Cannulae placement in the ePet-cre and the DAT-ires-cre mice. **Figure 3–source data 1.** fig3_5HT.xlsx contains AUC from all experiments with ePet-cre mice; related to figure 3A-C figure supplement 1A-C. **Figure 3–source data 2.** fig3_DA.xlsx contains AUC from all experiments with DAT-ires-cre mice; related to figure 3D-F figure supplement 1D-F. **Figure 3–source data 3.** Fig3_stats.ipynb contains statistical analysis related to figure 3 and its supplements.

### The LH→DRN, LH→VTA, and LH→LHb pathways are activated by aversive cues

The LH→VTA and LH→LHb pathways encode aversive cues for active defensive behaviors (***Barbano et al., 2020***; ***de Jong et al., 2019***; ***Trusel et al., 2019***). To determine whether the LH→DRN pathway play a complementary role, we trained mice using an active avoidance task while simultaneously monitoring activity at LH terminals in the DRN, VTA and LHb. The mice learned to associate a conditioned stimulus (CS) predicting an upcoming mild foot shock that could be escaped or avoided by crossing from one compartment to the other of the two-compartment box (***Figure 4***A). The mice quickly learned to avoid foot shocks, with the mean escape latency decreasing and the fraction of avoidance increasing in the late phase of training (***Figure 4***B). Activity at the LH→DRN, LH→VTA, and LH→LHb pathways was monitored during the early and late stages of learning. Mobility scores were calculated using the DeepLabCut software package for markerless animal pose estimations (***Mathis et al., 2018***). Our results show that activity following the aversive cue during late training increases significantly in all three LH pathways, suggesting that learning signals are encoded in these pathways. Activity increased further at the onset of the foot shock and the escape response during early and late training (***Figure 4***C). We then investigated activity in trials where avoidance occurred, with the mice avoiding the foot shock by shuttling to the other compartment of the two-compartment box before receiving the foot shock. We observed a significant increase in peri-event activity at onset of avoidance in all three pathways, which provides further support for their role in engaging self-motivated responses (***Figure 4***D), reminiscent of what we observed with the TST.

**Figure 4.**
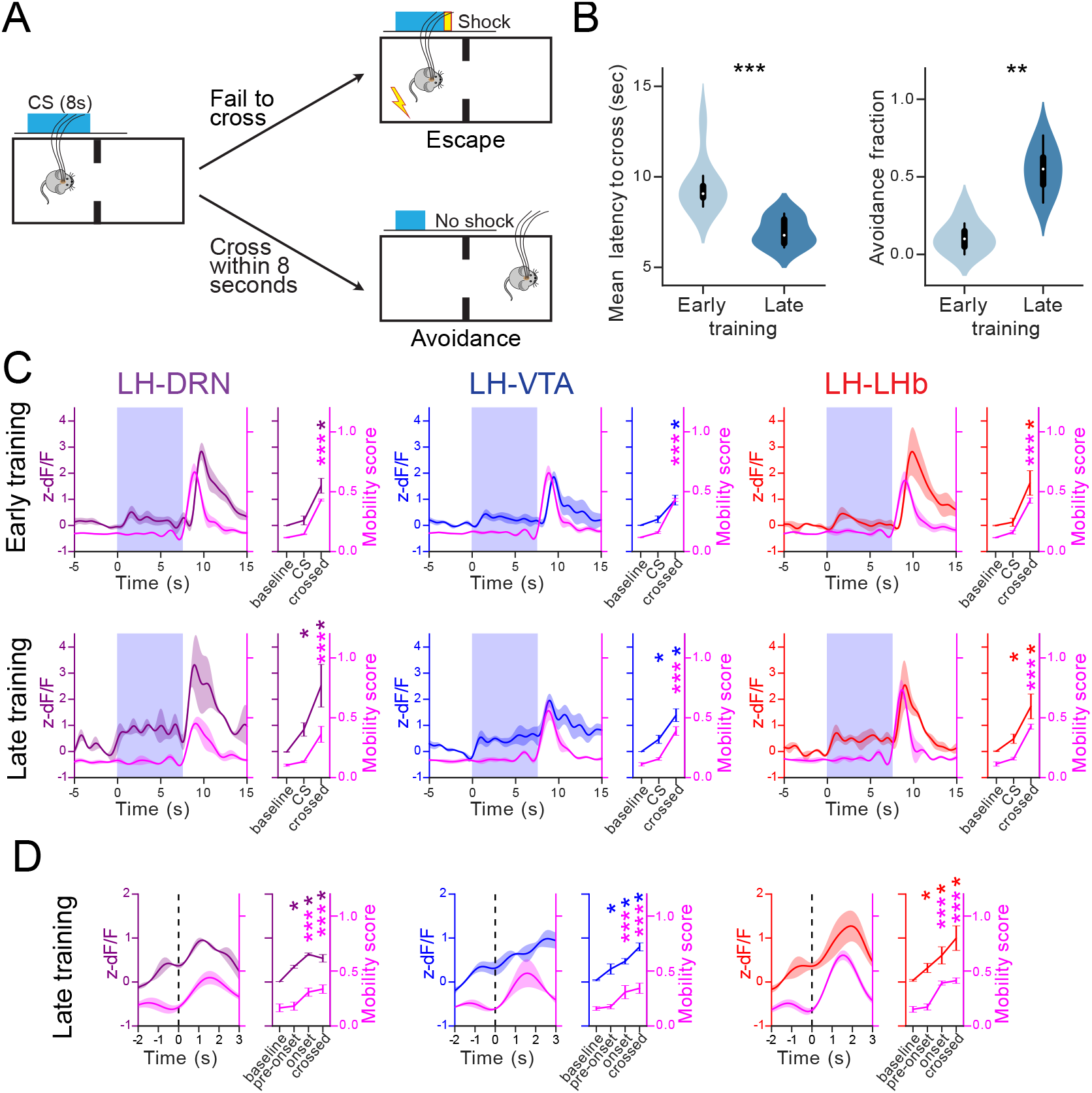
The LH→DRN, LH→VTA, and LH→LHb pathways are activated by aversive cues. (**A**) Diagram of the avoidance conditioning test. The mice learn to shuttle within 8 second of conditioned stimuli (CS, tone and light) in order to avoid a foot-shock. (**B**) The difference between early and late training trials is represented by the latency in shuttling from one compartment of the box to the other and the fraction of avoidance. (**C**) Peri-event plots of average *Ca*^2+^ signals to all CS events followed by escape at the LH→DRN, LH→VTA, and LH→LHb axonal terminals and plots for AUC at baseline, during CS and escape. The lines represent means ±SEM. The **top** plots represent the results during early training and the **bottom** plots represents the results during late training. Same convention as **C** for mobility onsets of avoidance during late training **D**. The magenta lines are mobility scores. Two-way repeated measures ANOVA within factors pathways (LH→DRN, LH→VTA, and LH→LHb) and time periods (different for CS and mobility onset) with post hoc Dunnett’s test. The *p* values were adjusted using the Bonferroni multiple testing correction method. \**p* < 0.05, \*\**p* < 0.01, \*\*\**p* < 0.001. **Figure 4–Figure supplement 1.** Cannulae placement. **Figure 4–source data 1.** fig4_LH_avoidanceTask.xlsx contains AUC from avoidance task with GCaMP6s-expressing mice; related to figure 4. **Figure 4–source data 2.** Fig4_stats.ipynb contains statistical analysis related to figure 4.

**Figure 5.**
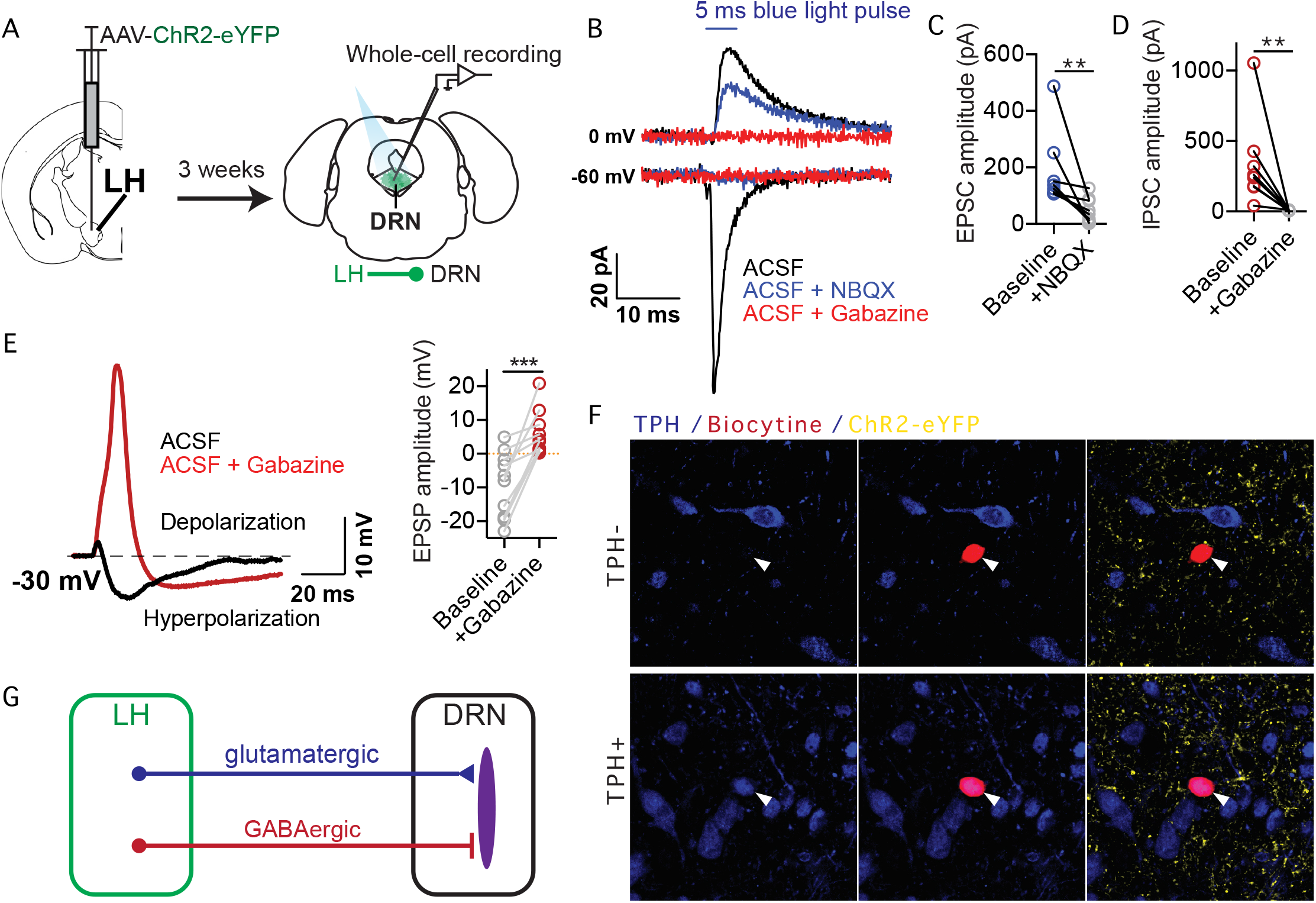
The LH provides monosynaptic excitatory and inhibitory projections to serotoninergic and non-serotoninergic neurons in the DRN. (**A**) Diagram of the experimental setup. (**B**) Representative monosynaptic response from a DRN neuron following optogenetic stimulation of LH axon terminals. The response involves both an excitatory inward current measured at −60 mV and an inhibitory outward current measured at 0 mV, which were respectively blocked by the AMPA receptor antagonist (NBQX, blue) and the GABA-a receptor antagonist (gabazine, red). Plots of ChR2-evoked EPSCs (**C**) and IPSCs (**D**) before and after blockade with the respective antagonists. (**E**) Representative monosynaptic response from a DRN neuron at a −30 mV holding voltage following optogenetic stimulation of LH axon terminals (left), and plots of EPSP amplitudes before and after GABA-a receptor blockade with gabazine (right). Gabazine abolished the hyperpolarizing inhibitory component, resulting in large excitatory depolarizing responses. (**F**) Confocal images of DRN neurons filled with biocytin (red) during whole cell patch-clamp recordings and post hoc immuno-labelling for the serotoninergic marker tryptophane hydroxylase (TPH, blue). LH axon terminals are shown in yellow in the DRN expressing ChR2-eYFP. (**G**) Schematic of connectivity between LH and DRN. Paired Wilcoxon test.\*\**p* < 0.01, \*\*\**p* < 0.001. **Figure 5–source data 1.** fig5_ephys.xlsx contains EPSC and IPSC amplitudes; related to figure 5. **Figure 5–source data 2.** Fig5_stats.ipynb contains statistical analysis related to figure 5.

### The LH provides monosynaptic excitatory and inhibitory inputs to serotoninergic and non-serotoninergic neurons in the DRN

Previous studies have shown that LH neurons control the LHb through excitatory inputs while sending both excitatory and inhibitory projections to the VTA, differentially targeting dopaminergic and GABAergic neurons (***de Jong et al., 2019***; ***Barbano et al., 2016***, ***2020***; ***Nieh et al., 2016***). To examine synaptic transmission from the LH to DRN, the LH of wild-type mice were injected with AAV-ChR2-eYFP. Three weeks later, acute brain slices encompassing the DRN were prepared, and whole-cell responses were obtained from DRN neurons. All the recordings were made in the presence of TTX and 4-AP to avoid polysynaptic responses. Recorded neurons were filled with biocytin and were identified as serotoninergic or non-serotoninergic by post hoc staining for tryptophane hydroxylase (TPH). In voltage-clamp mode recordings, 5-ms light pulses evoked excitatory inward currents and inhibitory outward currents at −60 mV and 0 mV holding currents, respectively. Monosynaptic excitatory and inhibitory transmissions were blocked with the AMPA receptor antagonist NBQX and the GABA-a receptor antagonist gabazine, respectively. In whole-cell current clamp mode recordings with a −30 mV depolarized voltage and in presence of TTX and 4-AP, single light pulse induced a small depolarization followed by a large hyperpolarization. In the presence of gabazine however, the single light-pulses revealed a large excitatory depolarizing component. No major difference were observed between serotoninergic and non-serotoninergic DRN neurons (not shown). These results confirm that LH sends convergent monosynaptic excitatory and inhibitory inputs to the DRN (***Zhou et al., 2017***), and suggest that LH inhibitory transmission gate excitatory transmission on both serotoninergic and non-serotoninergic neurons in the DRN.

### Optostimulation of the LH→DRN, LH→VTA, and LH→LHb pathways increases mobility

To determine whether increased activity at LH neural outputs is sufficient to motivate behavior, the LH of wild-type mice were injected with an AAV encoding the opsin Channelrhodopsine-2 (ChR2) fused to the fluorescent protein eYFP (AAV-ChR2-eYFP), or with eYFP alone (AAV-eYFP) as a control. Optical fibers were implanted as described for the fiber photometry recordings (***Figure 6***A). A single optical fiber cannula was connected to a 450-nm laser light source for the optogenetic stimulation of individual LH neural outputs. All three LH neural outputs were tested using a Latin square experimental design, with 24 h between experimental sessions. The mice were subjected to 20-min TST sessions. Each session consisted of 2-min epochs with or without 5-ms light pulses delivered in 20Hz optical stimulation trains (***Figure 6***B). The stimulation of individual LH neural outputs was sufficient to increase mobility during the 2-min stimulation epochs compared to the non-stimulation epochs in the TST (***Figure 6***C). The stimulation had no effect on the mobility of mice expressing eYFP only (***Figure 6***D). The same results were also obtained in the OFT (***Figure 6–Figure Supplement 1***). These results show that increased activity at individual LH neural outputs is sufficient to promote motivated behavior.

**Figure 6.**
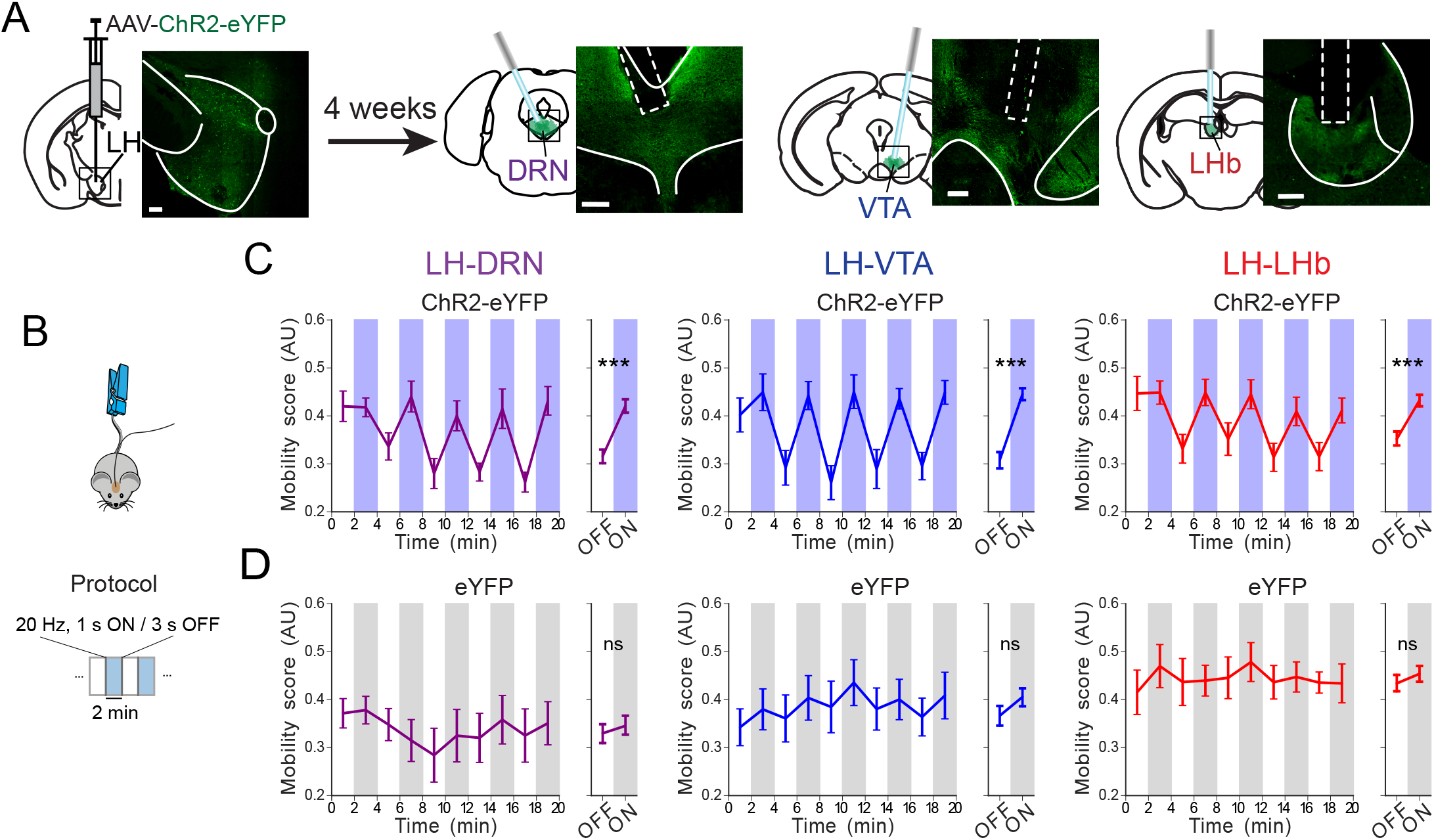
Optostimulation of the LH→DRN, LH→VTA, or LH→LHb pathway increases active coping behavior in the TST. (**A**) Diagram of the experimental setup and fluorescence images of ChR2-eYFP expression in LH neurons and axon terminals at the DRN, VTA, and LHb (scale bar 200 *μ*m). (**B**) Diagram of the experimental protocol for the TST. The mobility of the mice was evaluated during a 20 min TST session with alternating 2-min epochs without or with 1-s, 20 Hz trains every 4 sec. Mobility was automatically monitored with a video tracking system. (**C-D**) Plots of mean mobility (mean ± SEM) during periods of optogenetic stimulation (blue (**C**) or gray (**D**)) or no light (white) at the LH→DRN (**left**), LH→VTA (**middle**), or LHA→LHb (**right**) pathways in ChR2-eYFP- (**C**) and eYFP-expressing (**D**) mice. Four-way repeated measures ANOVA between factors of group (ChR2-eYFP- and eYFP-expressing mice), and within factors pathway (LH→DRN, LH→VTA, or LH→LHb), time period (five 4-minutes periods) and laser (on and off) with post hoc Tukey test. The *p* values were adjusted using the Bonferroni multiple testing correction method. \*\*\**p* < 0.001, *p* > 0.2 ns (not significant). **Figure 6–Figure supplement 1.** Optostimulation of the LH→DRN, LH→VTA, or LH→LHb pathway increases mobility in the OFT. **Figure 6–Figure supplement 2.** Cannulae placement in mice expressing ChR2-eYFP **Figure 6–Figure supplement 3.** Cannulae placement in mice expressing eYFP **Figure 6–source data 1.** fig6_LHopto_TST-OFT.xlsx contains results of analysis of experiments in the TST and the OFT for mice expressing ChR2-eYFP and eYFP; related to figure 6 and figure supplement 1. **Figure 6–source data 2.** Fig6_stats.ipynb contains statistical analysis related to figure 6.

### Optostimulation of LH neural outputs has distinct effects in place preference and sucrose consumption

Consistent with previous studies, we found that the stimulation of the LH→LHb pathway was aversive. The mice avoided a context paired with the stimulation during a real-time place preference (RTPP) test (***Figure 7–Figure Supplement 1***A-B). However, even though the stimulation of the LH→DRN and LH→VTA pathways significantly increased mobility in the TST and OFT, it did not promote place preference or place avoidance in the RTPP. To further investigate the contribution of LH neural outputs in reward-related behaviors, we examined the effect of optogenetic stimulation on sucrose consumption (***Figure 7***). To this end, water-deprived mice were allowed to drink the sucrose solution as described in the fiber photometry experiments, and 1-s, 20-Hz optogenetic stimulation were delivered at the onset of drinking events (***Figure 7***A). The effect of the optogenetic stimulation was compared to sessions without optogenetic manipulation (see Methods). The optogenetic stimulation of the LH→LHb pathway at the onset of drinking events decreased average drinking duration and total consumption, which is consistent with previous findings (***Stamatakis et al., 2016***). Conversely, optogenetic stimulation of the LH→VTA and LH→DRN pathways did not decrease drinking duration, but rather, decreased the time between drinking events, increased the total number of drinking events, and consequently increased total sucrose consumption (***Figure 7***B-E). Taken together, these results suggest that LH neural outputs use different mechanisms to control motivated behaviors.

**Figure 7.**
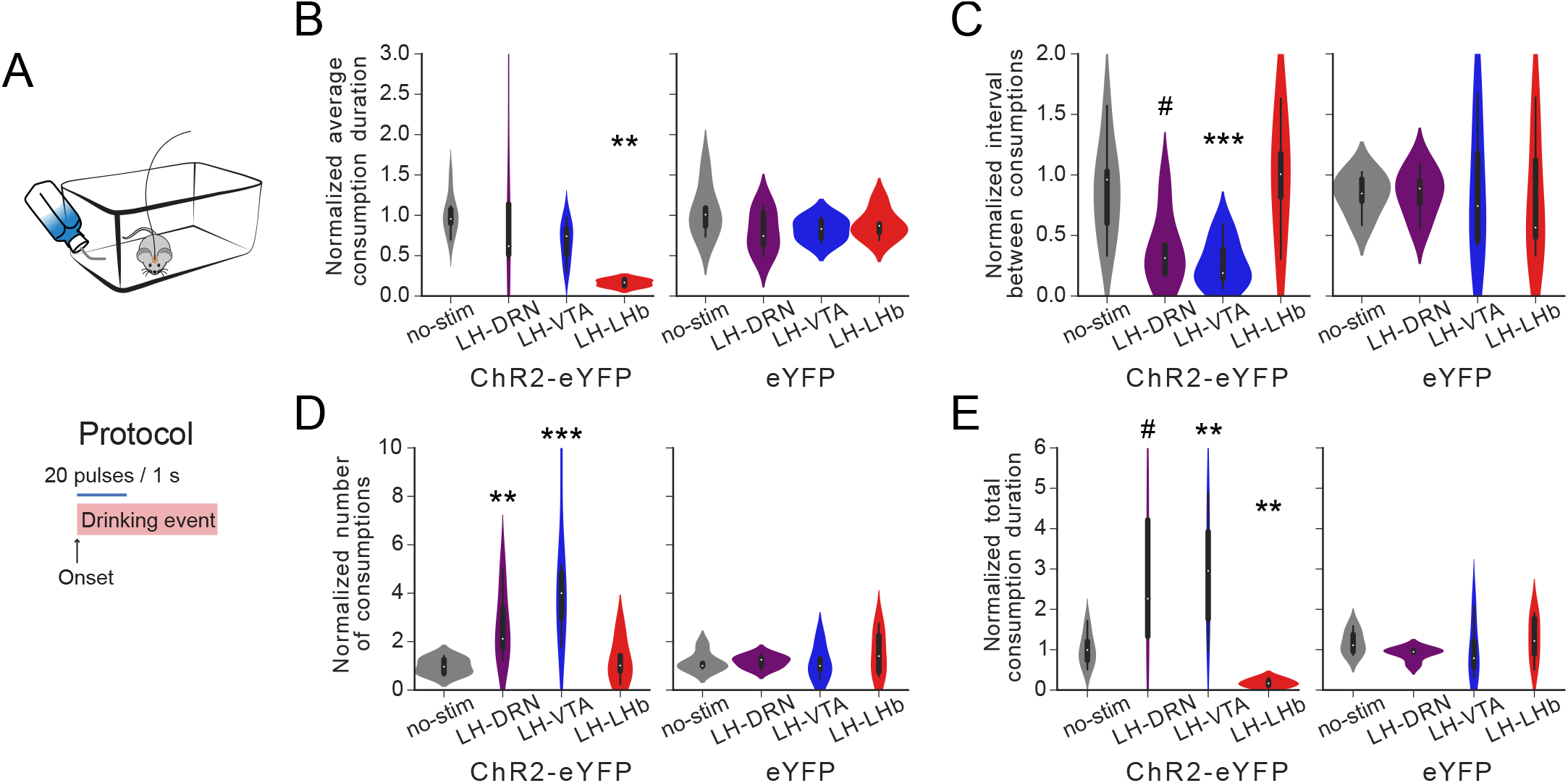
Optostimulation of LH neural outputs had distinct effects on sucrose consumption. (**A**) Diagram of the experimental protocol during sucrose consumption. A 1-s, 20 Hz optogenetic stimulation was given at the onset of a drinking event. (**B-E**) Plots of normalized average drinking duration (**B**), average interval between consumption events (**C**), total number of consumption events (**D**), and total drinking duration (**E**) normalized to a session without optogenetic stimulation for experimental mice expressing ChR2-eYFP (left panels) and control mice expressing eYFP only (right panels). Two-way ANOVA between factor of group (ChR2-eYFP- and eYFP-expressing mice) and within factor stimulated pathway (no stimulation, LH→DRN, LH→VTA, or LH→LHb) with post hoc Dunnett’s. The *p* values were adjusted using the Bonferroni multiple testing correction method. #*p* < 0.1,\**p* < 0.05, \*\**p* < 0.01, \*\*\**p* < 0.001, *p* > 0.2 ns (not significant). **Figure 7–Figure supplement 1.** Optostimulation of the LH→DRN, LH→VTA, or LH→LHb pathways in the RTPP. **Figure 7–source data 1.** fig7_LHopto_RTPP-SCT.xlsx contains results of analysis of experiments in the RTPP and the SCT for mice expressing ChR2-eYFP and eYFP; related to figure 7 and figure supplement 1. **Figure 7–source data 2.** Fig7_stats.ipynb contains statistical analysis related to figure 7.

## Discussion

The LH is a central nucleus that connects many brain regions that orchestrates vital behaviours (***Bonnavion et al., 2016***; ***Fakhoury et al., 2020***). It is the central brain region of an interconnected network that includes the DRN, VTA and the LHb, which all have important functions in reward and sensory processing, arousal, and motivation (***Bonnavion et al., 2016***; ***Proulx et al., 2014***; ***Nakamura, 2013***; ***Morales and Margolis, 2017***). However, most studies have focused on distinct neurochemically defined LH populations or single downstream projections (***Bonnavion et al., 2016***; ***Fakhoury et al., 2020***; ***de Jong et al., 2019***; ***Barbano et al., 2020***; ***Lecca et al., 2020***; ***Jennings et al., 2015***; ***Lazaridis et al., 2019***). No studies have explored how multiple LH neural inputs or outputs are dynamically recruited in live animals performing behavioral tasks. Here, we examined how three major LH neural outputs are dynamically engaged in tasks involving cue processing and motivated behavioral responses. We used multi-fiber and dual-color photometry recordings to show that there is significant coherence in neural activity at LH neural outputs projecting to the DRN, VTA and LHb. These projections are from largely non-overlapping LH neuronal populations. Increased activity at these three pathways preceded mobility onset in the TST and preceded changes in the activity of serotoninergic and dopaminergic neurons in the DRN and VTA, respectively. While the optogenetic stimulation of individual LH neural outputs was sufficient to increase the mobility of mice in the OFT and TST, it had a different effect on sucrose consumption. Optogenetic stimulation of the LH→LHb pathway reduced drinking duration and total sucrose consumption, whereas optogenetic stimulation of the LH→DRN and LH→VTA pathways decreased the interval between drinking events and, consequently increased the total number of drinking events and total sucrose consumption. These results indicates that these three LH neural outputs play complementary roles in engaging motivated behaviors.

Recent studies that measured neuronal activity in multiple brain regions have shown that coordinated neural network dynamics are important for organizing complex behaviors (***Allen et al., 2019***; ***Steinmetz et al., 2019***). In line with these findings, our results provide evidence that three LH neural outputs act in concert to engage motivated behavioral responses. Our results support the role of LH as a central hub for signaling aversive cues to the DRN, the VTA, and the LHb in learned and innate defensive behaviors (***Barbano et al., 2020***; ***de Jong et al., 2019***; ***Trusel et al., 2019***). Moreover, we also demonstrate that all three LH neural outputs play an important role in initiating cue-independent spontaneously motivated active coping responses.

We combined retrograde tracing and intersectional viral approaches to shown that only a few LH neurons project into two or more downstream brain regions. Our results further show that there are only a few collaterals, which is consistent with previous observations (***de Jong et al., 2019***). Our dual-color photometry recordings also confirmed that neural activity from two distinct LH neuronal populations projecting to DRN and VTA exhibit significant coherence. Taken together, these results are important for two reasons: (1) they show that independent LH neural populations send coherent signals to downstream brain regions, and (2) they support the validity of recording activity at axon terminals to study signal transmission along specific neural pathways.

The LHb receives a net excitatory input from the LH (***Lazaridis et al., 2019***; ***Trusel et al., 2019***). Recent work has shown that aversive stimuli are sufficient to activate the LH→LHb pathway and promote escape behavior (***Lecca et al., 2017***). Moreover, this pathway encodes negative valences and rapidly develops prediction signals for negative events and aversive cues, making this pathway a critical node for value processing and avoidance learning (***Lazaridis et al., 2019***; ***Trusel et al., 2019***). The results we obtained in our active avoidance task are consistent with these findings.

The VTA receives both excitatory and inhibitory inputs from the LH, both of which play roles in motivated behaviors (***Nieh et al., 2015***, ***2016***; ***Barbano et al., 2016***, ***2020***; ***de Jong et al., 2019***). Glutamatergic transmission at the LH→VTA pathway (LH^*Glut*^ →VTA) plays a role in motivating innate escape responses, signaling unexpected aversive outcomes, and signaling cues predicting aversive outcomes (***de Jong et al., 2019***; ***Barbano et al., 2020***). This aversive signal is mediated by the activation of VTA dopaminergic neurons projecting to the ventromedial section of the nucleus accumbens (***de Jong et al., 2019***). ***Barbano et al.*** (***2020***) showed that LH^*Glut*^ →VTA transmission mediates innate escape responses by the activation of VTA glutamatergic neurons. GABAergic transmission at the LH→VTA pathway (LH^*GABA*^ →VTA) plays an important role in promoting behavioral activation leading to compulsive sucrose-seeking and feeding (***Nieh et al., 2015***; ***Barbano et al., 2016***), behaviors that are mediated by the disinhibition of VTA dopamine neurons (***Nieh et al., 2015***). In light of these findings, it is likely that the increased activity observed in aversive contexts (AP, TST, active avoidance) is mediated by the LH^*Glut*^ →VTA transmission while the increased in activity measured prior to sucrose consumption is mediated by the LH^*GABA*^ →VTA transmission. Taken together, these results suggest that behavioral responses are engaged by an increase in the activity of DA neurons in the VTA through direct activation (aversive contexts) or disinhibition (reward-seeking behavior) of DA activity that originates from LH projections.

We show that the LH→DRN pathway is also involved in engaging motivated behavioral responses in freely behaving mice. Serotoninergic DRN neurons play important roles in organizing motivated and reward-related behaviors (***Nakamura, 2013***), in emotions and aversive processing (***Cools et al., 2008***), and in active coping behaviors (***Nishitani et al., 2019***; ***Cools et al., 2008***; ***Nakamura, 2013***). Here we show that the LH→DRN pathway is activated when mice are presented with an aversive airpuff or a mild foot shock. This pathway is also prominently engaged before the mice reach the reward spout, as we observed with the LH→VTA pathway and at mobility onset, which indicates that there is a significant correlation with mobility in the TST.

Both DRN^5*HT*^ and VTA^*DA*^ neurons increased their activity at the onset of mobility in coping behaviors such as the TST. This change in activity was preceded by the activation of the LH→DRN and LH→VTA pathways, suggesting that signal transmission from the LH leads to the activation of the DRN^5*HT*^ and VTA^*DA*^ neurons in order to control motivated responses.

In the DRN, serotoninergic and GABAergic neurons receive prominent inputs from the LH (***Ogawa and Watabe-Uchida, 2018***; ***Weissbourd et al., 2014***; ***Pollak Dorocic et al., 2014***; ***Celada et al., 2002***). ***Zhou et al.*** (***2017***) recently shown that functional excitatory and inhibitory inputs converge onto serotoninergic DRN neurons. Our *ex vivo* electrophysiological recordings showed similar inputs converging onto non-serotoninergic DRN neurons indicating that there is a complex functional relationship between the LH and the DRN (***Celada et al., 2002***). However, it remains to be determined whether this increase in serotoninergic activity is mediated by direct activation of 5-HT DRN neurons or by indirect disinhibition through local GABAergic neurons.

We also show that all three LH neural outputs are activated prior to the onset of movement in the TST, an energetically-demanding and aversive context (***Proulx et al., 2014***; ***Warden et al., 2012***). Recent miniscopic *in vivo* calcium imaging of LHb neurons in mice responding in looming experiments has revealed that specific LHb neuron ensembles are active before the mice start running away from an aversive threat (***Lecca et al., 2020***). Our results suggest that some of the excitatory inputs driving these neuronal ensembles may be provided by the LH. Our results are also reminiscent of recent studies that directly measured the activity of orexin-expressing LH neurons and shown that this cell population is activated by aversive stimuli and stressors and is inhibited during food consumption, which is independent of food taste and texture (***González et al., 2016a***,b). Moreover, the same group recently showed that a large fraction of orexin-expressing LH is activated at the onset of the initiation of movement (***Karnani et al., 2020***). We propose that signal transmission from this genetically-defined population in the LH play an important role in motivating the initiation of movement by processing reward and sensory information in the DRN, VTA, and LHb (***Flanigan et al., 2020***; ***Harris et al., 2005***). Since most orexin-expressing neurons in the LH corelease glutamate (***Rosin et al., 2003***; ***Mickelsen et al., 2019***), it would be interesting to determine how excitatory and orexigenic transmissions drive activity in the DRN, VTA, and LHb and as a result, control motivated behaviors.

In the SCT, there was also a significant increase in the activity at the LH→DRN and LH→VTA pathways prior to the mice reaching the sucrose dispenser, while no significant change in the activity at the LH→LHb pathway was observed in this task. These results suggest that the LH→DRN and LH→VTA pathways may play a more general role in motivating goal-directed behavior, while the LH→LHb pathway would be critically involved in orchestrating behavioral responses in aversive contexts. This notion is supported by previous experiments showing that the inhibition of the LH→LHb pathway impaired aversion-driven escape behavior (***Lecca et al., 2017***) and disrupted avoidance learning. The inhibition of the LH→VTA pathway also impaired innate defensive behavior (***Barbano et al., 2020***). Our optogenetic experiments also support this scenario. Although the optogenetic stimulation of individual LH neural outputs increased mobility in the TST, optogenetic stimulation at drinking onset in the SCT had a different effect. Indeed, the activation of the the LH→LHb pathway at drinking onset decreased drinking duration and total sucrose consumption, whereas the activation of the LH→DRN or the LH→VTA pathway increased total sucrose consumption by decreasing the time interval between drinking periods. One possible underlying mechanism that enhances sucrose consumption following LH→VTA activation may be mediated by the co-release of neurotensin from LH axon terminals in the VTA, a neuropeptide that promotes reward-seeking by enhancing glutamate transmission in the VTA (***Kempadoo et al., 2013***). In the SCT, phasic activity at the LH^*GABA*^ →VTA pathway at the onset of sucrose consumption may also be sufficient to motivate reward-seeking behaviors by decreasing time the interval between drinking periods (***Barbano et al., 2016***; ***Jennings et al., 2015***).

In summary, we show the LH→LHb, LH→VTA, and the LH→DRN pathways are important in active coping and defensive behaviours. For many contexts and stimuli, the responses of the LH→DRN, LH→VTA, and LH→LHb pathways are coherent, suggesting that the LH conveys complementary information that engages several downstream brain regions to elicit motivated behaviors. It is likely that distinct 5-HT DRN, and DA VTA neuronal populations, segregated by their inputs and outputs, play different but complementary roles in cue processing and adaptive behaviors. Another interesting question that remains to be investigated is whether the three distinct LH neuronal populations, independently projecting to DRN, VTA and LHb, receive common or distinct inputs from upstream brain regions.

## Methods and Materials

### Animals

All experiments were performed with 8− to 12-week-old wild-type C57Bl/6J, ePet-cre, or DAT-irescre mice. The mice were housed 2-4 per cage and were kept on a 12 h/12 h light/dark cycle. All the experiments were performed in accordance with the Canadian Guide for the Care and Use of Laboratory Animals guidelines and were approved by the Université Laval Animal Protection Committee.

### Stereotactic injections

The mice were anesthetized with isoflurane for the stereotaxic injections of adeno-associated viruses (AAVs) and the implantation of optical fibers. Viral titers ranged from 10^12^ to 10^13^ genome copies per milliliter and volumes ranged from 100 to 200 nl per side. Injections were performed using a glass pipette mounted on a stereotactic table. The AAVs were infused at a rate of 1 nl/sec. At the end of the injection, the pipet was left *in situ* for 5 min to allow the virus to diffuse into the surrounding tissue. Three to four weeks later, for the fiber photometry and optogenetic experiments, the mice were implanted with 200- or 400 *μ*m cannulae with a metal ferrule for the fiber photometry recordings, and 200 *μ*m cannulae with a ceramic ferrule for the optogenetic manipulations.

For most fiber photometry experiments, the mice were bilaterally injected with AAVDJ-CAG-GCaMP6s in the LH, and optical fiber cannulae were implanted with their tips immediately above the DRN, VTA, and LHb. For the intersectional viral strategy, the mice were injected with retroAAV-CAG-cre-P2A-mNeptune in the VTA, retroAAV-CAG-flpO-P2A-TagBFP2 in the DRN, and a mix of the AAV9-CAG-DIO-jrGECO and AAV9-EF1a-fDIO-GCaMP6s in the LH, and an optiocal fiber was chronically implanted above the LH. For the dual-color fiber photometry, ePet-cre mice were injected with AAVDJ-CAG-GCaMP6s in the LH and AAV9-CAG-DIO-jrGECO1a in the DRN, and optical fiber was implanted above the DRN. Similarly, DAT-ires-cre mice were injected with AAVDJ-CAG-GCaMP6s in the LH and AAV9-CAG-DIO-jrGECO1a in the VTA, and optical fiber was implanted above the VTA.

For the optogenetic experiments, the mice were bilaterally injected with AAV9-hSyn-hChR2(H134R)-eYFP or AAV9-hSyn-eYFP in the LH, and optical fibers were implanted above the DRN, VTA, and LHb. The mice were tested 2 weeks post-cannulae implantation to give them time to recover from the surgical procedures.

For the electrophysiology experiments, the mice were injected with AAV9-hSyn-hChR2(H134R)-eYFP in the LH. Acute brain slices were prepared 3 weeks later for whole-cell patch clamp recordings.

To label DRN-, VTA-, and LHb-projecting neurons in the LH of the same animal, the mice were injected with CTx488 (Alexa Fluor 488-conjugated cholera toxin subunit B), CTx594, and CTx647 in the DRN, VTA, and LHb, respectively. The mice were transcardially perfused 3 days later, and their brains were processed for histology.

The coordinates for the injections were as follows: LH : −1.2 mm AP, ±1.0 mm ML, −5.2 mm DV; LHb : −1.65 mm AP, ±0.45 mm ML, −2.8 mm DV; VTA : −3.3 mm AP, ±0.5 mm ML, −4.8 mm DV; DRN: −4.65 mm AP, 0.0 mm ML, −3.2 mm DV. The coordinates for implantation were the following: LH : −1.2 mm AP, −1.0 mm ML, −5.0 mm DV; LHb : −1.65 mm AP, −0.45 mm ML, −2.4 mm DV; VTA : −3.3 mm AP, 1.0 mm ML, −4.4 mm DV ∠10°; DRN : −4.65 mm AP, −1.05 mm ML, −2.9 mm DV ∠20°.

### Fiber photometry recordings

A custom-build (***Kim et al., 2016***) or a Neurophotometrics (https://neurophotometrics.com) fiber photometry systems was used to record calcium signals. Both systems had the same set of dichroic mirrors, filters, and LEDs. Light from 415-nm, 470-nm, and 560-nm LEDs were bandpass filtered, collimated, reflected by dichroic mirrors, and focused by a 20× objective (numerical aperture, NA 0.39). The light passed through a patch cord of three fibers, that were connected to the implanted cannulae. The emitted fluorescence was collected by the same fibers, filtered, and separated into red and green images, which were projected on a CMOS camera sensor. The excitation power was adjusted so as to get 50 to 70 uW of each of the lights at the tip of the patch cord. The custom-build system was controlled using LabJack and a custom-written Matlab code. The Neurophotometrics system was run by Bonsai open source software. For most of the experiments, light from 415-nm-and 470-nm LEDs was alternated such that the camera captured fluorescent excitation light from either the 415-nm or 470-nm LED. The camera captured images at 20 Hz. Signals from the two excitation wavelengths were sampled at 10 Hz. For the experiments with the ePet-cre and DAT-ires-cre mice, the 560-nm LED was alternated with the 470-nm LED. For the intersectional viral strategy experiments, all three (415-nm, 470-nm, and 560-nm) LEDs were alternated such that the camera captured one light at a time. See ***Martianova et al.*** (***2019***) for more details.

The mice were connected to a 3-fiber patch cord to record the signals during all the tests. For intersectional viral strategy experiments with the ePet-cre and DAT-ires-cre mice, only one fiber was connected.

#### Airpuff test

The mice were placed in the open field arena and airpuffs were delivered on top of the animals every 60 s, for a total of 5 airpuffs per 6 min session. Airpuff delivery was paired with a key press on a computer keyboard that was registered by the fiber photometry software and that provided timestamps of the airpuffs.

#### Sucrose consumption test

The mice were water-deprived for 24 h before the experiment. For the test, the mice were placed in a cage with free access to a spout delivering a small amount of sucrose solution. Consumption periods were automatically tracked with a custom lickometer. The setup to measure licks consisted of a mouse cage covered with a metal grid floor. The cage was equipped with a copper wire-wrapped metal sipper tube from which sucrose solution was delivered. Each lick closed an electrical circuit for the duration of the contact with the sipper tube. The junction potential between the metal sipper tube and the mouse was recorded using the ANY-maze system. ANY-maze provided timestamps of each of the loop closures. To align the lick timestamps with the fiber photometry recordings, the start of the ANY-maze test was either triggered by the fiber photometry system or by ANY-maze, and the fiber photometry software recorded the actual timestamps of the same computer.

#### Tail suspension test

The mice were suspended by their tail for 10 min. Movement was monitored using a camera and ANY-maze video-tracking software. ANY-maze detects the speed of animal movement. The ANYmaze recordings were aligned to the fiber photometry recordings using the same strategy as with the sucrose consumption test.

#### Open field test

The mice were placed in an open field (50 cm × 50 cm) and movement was tracked using a camera and ANY-maze video-tracking software as with to the tail-suspension test.

#### Avoidance Test

A mouse was placed in a two-compartment chamber (Med Associates). A conditioned stimulus (CS, light, and tone) was provided for 8 s pseudo-randomly with an average ITI of 40 s. At the end of the 8-s CS, a mild foot-shock (0.2–0.4 mA) was delivered through the grid floor for 8 s or until the mouse crossed to the other compartment, which stopped the shock. If the mouse crossed within the 8-s CS, no shock was delivered. This was referred as avoidance. The mice were tracked using a camera and ANY-maze video-tracking software. To align the recordings of all the setups, Med Associates software triggered the start of the fiber photometry and the ANY-maze software at the beginning of the tests. The position of the mouse was analyzed post hoc using DeepLabCut, a toolbox for markerless animal pose estimation. (***Mathis et al., 2018***).

### Optogenetic manipulations

For the optogenetic experiments, the mice were connected to a 450-nm laser (Doric Lenses) through an optical fiber and a rotary joint. Pulses of blue light were controlled by the ANY-maze software. The stimulation protocol was 20-Hz trains and 5-ms pulses for 1 s every 4 s. The light intensity was adjusted to provide 10 mW at the tips of the implanted optical fiber cannulae. During a trial, the mice were connected to a single optical fiber cannula implanted above the DRN, VTA, or LHb. For each test, the mice were tested on consecutive days using a Latin square experimental design.

#### Real-time place preference test

The mice were placed in a chamber with two compartments connected by a small corridor. After 1 min of habituation, one of the compartments was paired with an optogenetic stimulation. A mouse received a photostimulation every time it entered the paired chamber (randomly assigned). To maximize novelty and exploratory behavior on consecutive testing days, the RTPP apparatus was used as follows: day 1 with a plain floor, day 2 with bedding covering the floor, and day three with finely ground food pellet on the floor. Stimulation chambers were randomly assigned on each of the three days of testing. The location of the mouse (chamber 1, chamber 2, or corridor) was tracked, and laser activation was controlled using the ANY-maze video-tracking system.

#### Tail suspension test and open field test

The mice were suspended by their tail for 20 min. The photostimulation was alternated between a 2-min periods without stimulation and with a 2-min period with photostimulation trains (20 Hz, 1 second, 5-ms pulse duration, every 4 seconds).

#### Sucrose consumption test

The same protocol used for the fiber photometry recordings was used for the optogenetic manipulations. The ANY-maze tracking system detected drinking onset and triggered 1-s 20-Hz laser photostimulations for each drinking event. To define the baseline of drinking behavior for each animal, the mice were tested 3 times (without photostimulation) prior to the photostimulation sessions, and 1 time after (without photostimulation). The DRN, VTA, and LHb photostimulation sessions and the sessions with no photostimulation were alternated using a Latin square experimental design.

### Electrophysiology

Three weeks after the injection of AAV-ChR2-eYFP in the LH, the mice were anesthetized with isoflurane and were perfused transcardially with 10 mL of ice-cold NMDG-artificial cerebrospinal fluid (aCSF) solution containing (in mM): 1.25 NaH2PO4, 2.5 KCl, 10 MgCl2, 20 HEPES, 0.5 CaCl2, 24 NaHCO3, 8 D-glucose, 5 L-ascorbate, 3 Na-pyruvate, 2 thiourea, and 93 NMDG (osmolarity was adjusted to 300–310 mOsmol/L with sucrose). The pH was adjusted to 7.4 using 10 N HCl. Kynurenic acid (2 mM) was added to the perfusion solution on the day of the experiment. The brains were then quickly removed, and 250 *μ*m acute brain slices encompassing the DRN were prepared using a Leica VT1200S vibratome. The slices were placed in a 32°C oxygenated perfusion solution for 10 min and were then incubated for 1 h at room temperature in HEPES-aCSF solution (in mM): 1.25 NaH2PO4, 2.5 KCl, 10 MgCl2, 20 HEPES, 0.5 CaCl2, 24 NaHCO3, 2.5 D-glucose, 5 L-ascorbate, 1 Na-pyruvate, 2 thiourea, 92 NaCl, and 20 sucrose (osmolarity was adjusted to 300–310 mOsmol/L with sucrose). The pH was adjusted to 7.4 using 10 N HCl. They were then transferred to a recording chamber on the stage of an upright microscope (Zeiss) where they were perfused with 3-4 mL/min of aCSF (in mM): 120 NaCl, 5 HEPES, 2.5 KCl, 1.2 NaH2P04, 2 MgCl2, 2 CaCl2, 2.5 glucose, 24 NaHCO3, and 7.5 sucrose). The perfusion chamber and the aCSF were kept at 32°C. All the solutions were oxygenated with 95% O2/5% CO2. A 60x water immersion objective and a video camera (Zeiss) were used to visualize neurons in the DRN. Borosilicate glass (3-7 MΩ resistance) recording pipettes were pulled using a P-1000 Flaming/ Brown micropipette puller (Sutter Instruments). Recordings were performed using an Axopatch 200B amplifier (Molecular Devices). For the voltage-clamp recordings, the intracellular solution consisted of (in mM): 115 cesium methanesulfonate, 20 cesium chloride, 10 HEPES, 2.5 MgCl2, 4 Na2ATP, 0.4 Na3GTP, 10 Na-phosphocreatine, 0.6 EGTA, and 5 QX314, as well as 0.2% biocytin (pH 7.35). For the current-clamp recordings, the intracellular solution consisted of (in mM): 130 K-gluconate, 5 KCl, 10 HEPES, 2.5 MgCl2, 4 Na2ATP, 0.4 Na3GTP, 10 Na-phosphocreatine, and 0.6 EGTA (pH 7.35). Signals were filtered at 5 kHz using a Digidata 1500A data acquisition interface (Molecular Devices, San Jose, CA) and acquired using pClamp 10.6 software (Molecular Devices). Pipette and cell capacitance were fully compensated. To examine monosynaptic transmission, the extracellular recording solution was supplemented with 1 *μ*M TTX and 100 μM 4-AP. For the voltage-clamp experiments, postsynaptic currents were measured in DRN neurons clamped at −60 mV and 0 mV holding voltage following optogenetic stimulation of LH axon terminals with 5-ms blue light pulses delivered through the objective with a Colibri 7 LED light source (Zeiss). Excitatory and inhibitory transmissions were blocked with 3 mM NBQX and 10 mM gabazine, which are AMPA and GABA-a receptor antagonists, respectively. For the current-clamp experiments, DRN neurons were depolarized at −30mV and changes in the postsynaptic potential were measured before and after the addition of 10 mM gabazine. Once the recordings were completed, the slices were fixed in 4% formaldehyde for 30 min and were then transferred to a 0.1M phosphate buffer solution for post hoc histological analysis.

### Histology and immunostaining

The mice were deeply anesthetized using a mix of ketamine/xylazine (100 and 10 mg/kg, respectively, intraperitoneally) and were transcardially perfused with saline followed by a 0.1 M phosphate buffer solution (PB, pH7.4) containing 4% paraformaldehyde. The brains were postfixed overnight in the same solution, rinsed with PB, and stored in PB. Brain sections (100 *μ*m for histology and 50 *μ*m for immunostaining) were cut with a vibratome along the coronal plane.

Sections used for fiber photometry recordings and optogenetic manipulations were examined to confirm injection sites and cannulae placements. Recordings were excluded post hoc in the rare cases where an optical fiber was misplaced or where the expression of the construct of interest was off-target or low.

DRN sections from ePet-cre mice and VTA sections from DAT-ires-cre mice were stained for TPH (tryptophan hydroxylase, a marker for 5HT neurons) and for TH (tyrosine hydroxylase, a marker for DA neurons), respectively, using a standard 2-day immunostaining protocol. Briefly, free-floating slices were first blocked in PB containing 10% normal donkey serum (NDS) and 0.2% Triton X-100 for 1 h. They were then incubated overnight with primary antibodies diluted in PB containing 3% NDS and 0.2% Triton X-100 and then with a secondary antibody diluted in PB containing 3% NDS for 2 h at RT. The primary antibodies were anti-TPH (Millipore, sheep polyclonal, 1:1000 dilution) and anti-TH (Millipore, sheep polyclonal, 1:1000 dilution). The secondary antibody was donkey anti-sheep IgG DyLight™ 405 (Jackson ImmunoResearch, 1:500 dilution).

Sections from the electrophysiological recordings were immunostained for TPH using the protocol descried above. Immunostained sections and sections from mice injected with CTx were mounted using Fluoromount™ Aqueous Mounting Medium (Millipore-Sigma) and imaged using a Zeiss LSM700 confocal microscope.

### Data analysis

#### Fiber photometry data

To store, process, analyze, and visualize the fiber photometry data, a Python package of objects and functions was created, which is available at Fiber Photometry Data Analysis GitHub repository.

To calculate the standardized *dF* /*F* (*z*-score *dF* /*F* , *zdF* /*F*), the algorithm described in ***Martianova et al.*** (***2019***) was used. Briefly, calcium-dependent (*signals*) and calcium-independent (*references*) traces were smoothed using a band-pass filter (moving average was used in ***Martianova et al.*** (***2019***)), flattened by removing the baseline using an airPLS algorithm (adaptive iteratively reweighted Penalized Least Squares (***Zhang et al., 2010***)), and standardized to a mean value of 0 and a standard deviation of 1. *References* were fitted to *signals* using a non-negative robust least squares regression (Lasso algorithm), and *zdF* /*F* was calculated by subtraction from standardized *signals* standardized and fitted *references*.

To define the onsets and offsets of consumption, lickings recorded with the lickometer were used: onsets were defined as licks that persisted for at least 0.5 s, and offsets defined as the absence of licks for at least 1 s.

For most of the tests, ANY-maze provided the speed of the mice over time. For the avoidance conditioning test, the coordinates of the mice over time were exported from videos using the DeepLabCut toolbox (***Mathis et al., 2018***), and the speed was calculated using the following formula: 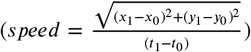. To account for different tracking setups and experimental setups, all the speeds were transformed to a standardized range [0, 1] (mobility score).

A mobility score was used to define mobility and immobility onsets. An immobility bout was defined as a period where the values were < 0.1 and lasted at least 2 s.

*zdF* /*F* was aligned and averaged around the specific events such as airpuff, consumption onset, mobility onset, CS, across all trials for each animal. To measure the change from a baseline area under the curve (AUC) was calculated. For airpuff, baseline AUC was calculated at −2 – −1 s, airpuff - at 0 – 1.5 s. For consumption onset, baseline AUC was calculated at −2 – −1 s, onset – at −0.5 – 1 s, drinking – at 2 – 4 s. For immobility onset, immobile AUC was calculated at −3 – −1 s, pre-onset – at −1 – 0 s, onset – at 0 – 1 s, mobile – at 1 – 3 s. In avoidance test, baseline AUC was calculated at −2 – −1 s, CS – at 0 – 8 s, crossed – at 8 – 11 s. AUC was normalized to the duration of the area.

#### Optogenetics data

For the RTPP, ANY-maze data of laser activity over time was exported from ANY-maze and the preference score was calculated using the following formula: 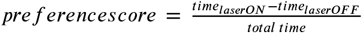 as detailed before (***Proulx et al., 2018***).

For the OFT and TST, the speeds tracked by ANY-maze were transformed into a standardized range [0, 1] (mobility score) to maintain the same format as the fiber photometry data. Average mobility scores were calculated for periods of laser ON and OFF.

For the SCT, ANY-maze data of licks recorded over time were exported, and the consumption onsets and offsets were found as described above. The following parameters were calculated: average consumption duration, average interval between events, number of events, and total consumption duration for three pre-sessions without photostimulation, sessions with photostimulation, and one post-session without stimulation. As the last pre-session and the post-session had approximately the same values for the parameters measured, these values were averaged and were used for the data normalization of values acquired during sessions with photostimulation.

#### Statistical analyses

Statistical analyses were performed using R programming language. Data distributions were tested using the Shapiro-Wilk normality test. Parametric or nonparametric tests were chosen depending on the number of observations, the distribution, and the model. For most of the data, a multi-factor mixed ANOVA with post hoc Dunnett’s or Tukey tests was used . The exact tests are indicated in the figure legends and ipynb source data files. Significance was set at p<0.05 in most tests. Electrophysiological and behavior experiments were replicated at least three times. Sample size were estimated from previously published work and from pilot experiments performed in our laboratory.

### Data and code availability

Raw data and the analysis steps from all the fiber photometry recordings and optogenetics manipulations were saved in hierarchical data format (h5) files. A Python code with the classes, functions, and an example of steps of analysis, including statistical analysis, was saved in Jupyter notebooks and is available at the following GitHub repository: Fiber Photometry Data Analysis

## Acknowledgments

We would like to thank Drs Armen Saghatelyan, Martin Lévesque, Sage Aronson and Sadegh Nabavi for their critical reviews of the manuscript. We also thank the Plateforme d’Outils Moléculaires at the CERVO Brain Center for producing the viral vectors, and Martin Lévesque’s lab for providing the DAT-ires-cre mice. CDP is supported by the Canadian Institutes of Health Research grant PJT169117 and the Natural Science and Engineering Research Council of Canada grant RGPIN-2017-06131, and receives Fonds de Recherche en Santé du Québec (FRQS) Junior-1 salary support. EM is supported by a doctoral training scholarship from the FRQS.

---

**Figure 1–Figure supplement 1.**
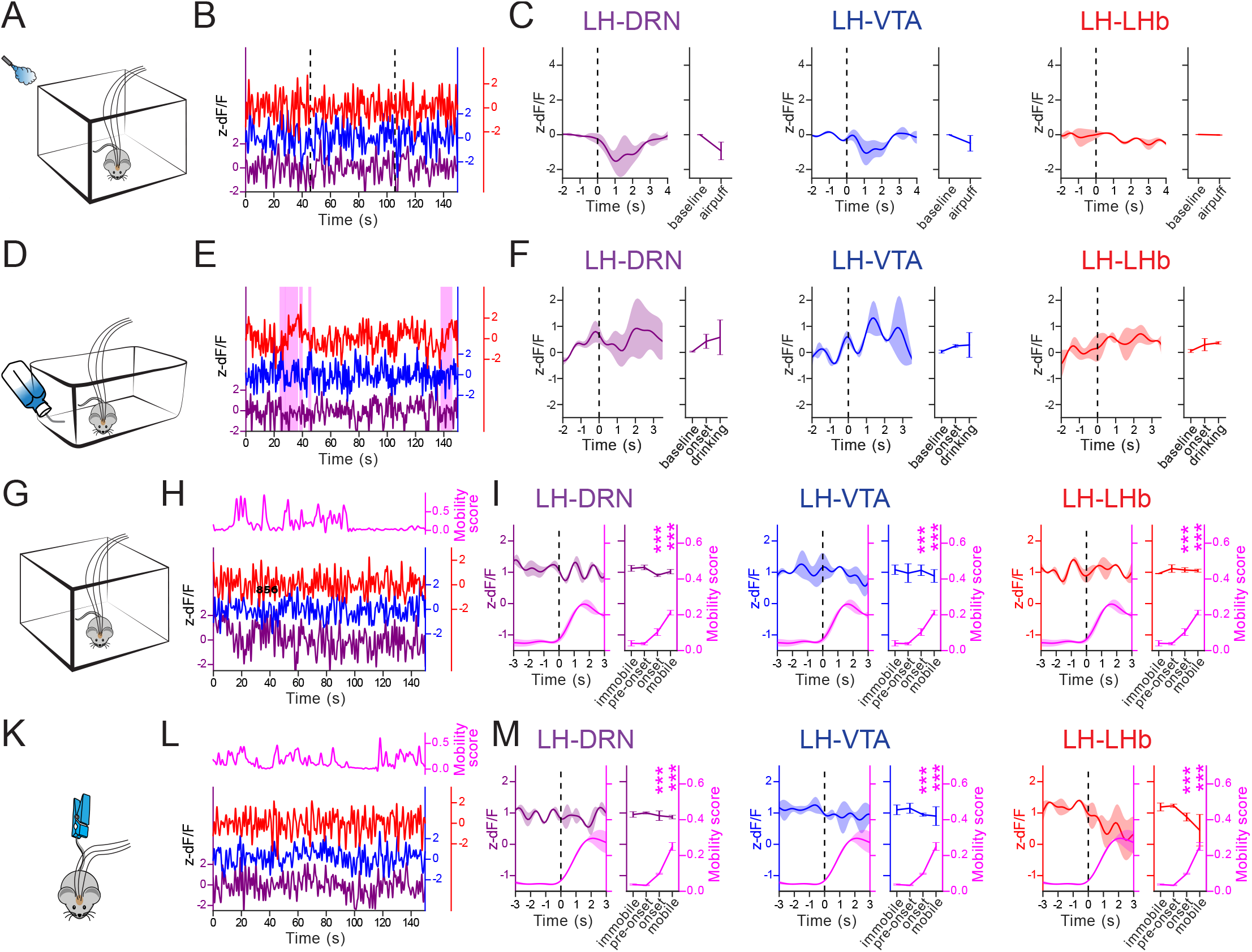
Recordings from the LH→DRN, LH→VTA, and LH→LHb pathways of control eYFP-expressing mice. (**A**) Diagram of the experimental setup for the airpuff. (**B**) Representative signal traces associated to the airpuffs (dashed vertical bars) simultaneously measured at the LH→DRN, LH→VTA, and LH→LHb pathways. (**C**) Peri-event plot of the average signals to all the airpuff events at the LH→DRN, LH→VTA, and LH→LHb pathways. Plot for average response before and after airpuffs. Lines represent mean ± SEM. Same convention as **D-F** for sucrose consumption test (**G-I**), open field test (**K-M**), and tail suspension test (**N-P**). The sucrose consumption events are represented by pink shaded box in **H**. The magenta lines are the mobility scores. The statistical analysis was performed along with the data from the mice expressing GCaMP6s.

**Figure 1–Figure supplement 2.**
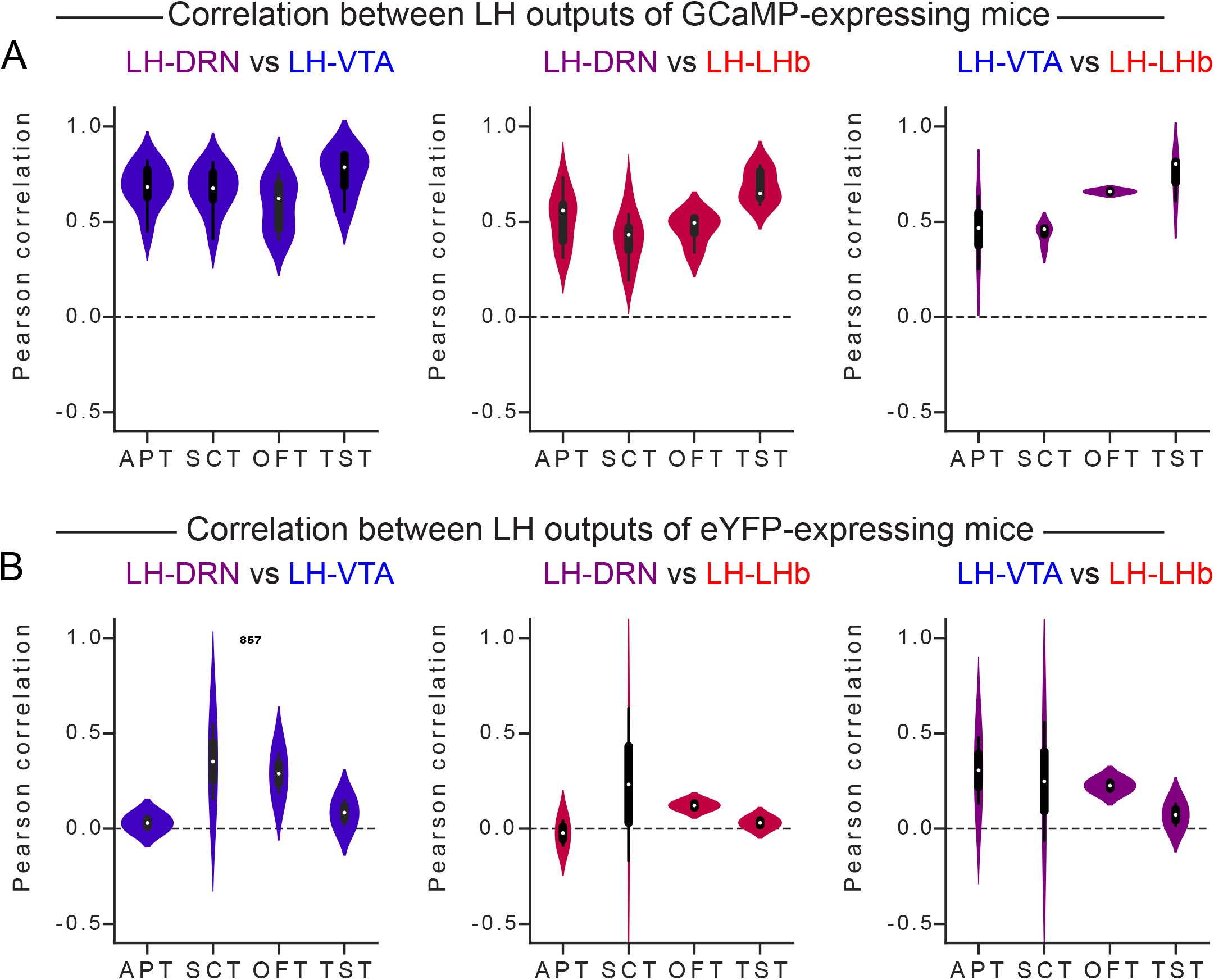
Pearson correlation between the signals recorded at the LH→DRN, LH→VTA, and LH→LHb pathways in mice expressing GCaMP6s and eYFP. (**A**) Pearson correlation between the *Ca*^2+^ signals recorded at the LH→DRN, LH→VTA, and LH→LHb pathways in mice expressing GCaMP6s. (**B**) Pearson correlation between the *Ca*^2+^ signals recorded at the LH→DRN, LH→VTA, and LH→LHb pathways in mice expressing eYFP. Three-way ANOVA between factors group (GCaMP6s- and eYFP-expressing mice), and within factors pathway (LH→DRN, LH→VTA, and LH→LHb), and tests (APT, airpuff test; SCT, sucrose consumption test; OFT, open field test; TST, tail suspension test) with post hoc Tukey test. The *p* values were adjusted using the Bonferroni multiple testing correction method. The main effect was the difference between the mice expressing GCaMP6s and eYFP (*p* < 0.05). One sample t-test showed significant difference from 0 in mice expressing GCaMP6s (*p* < 0.05), but not in mice expressing eYFP (*p* > 0.2)

**Figure 1–Figure supplement 3.**
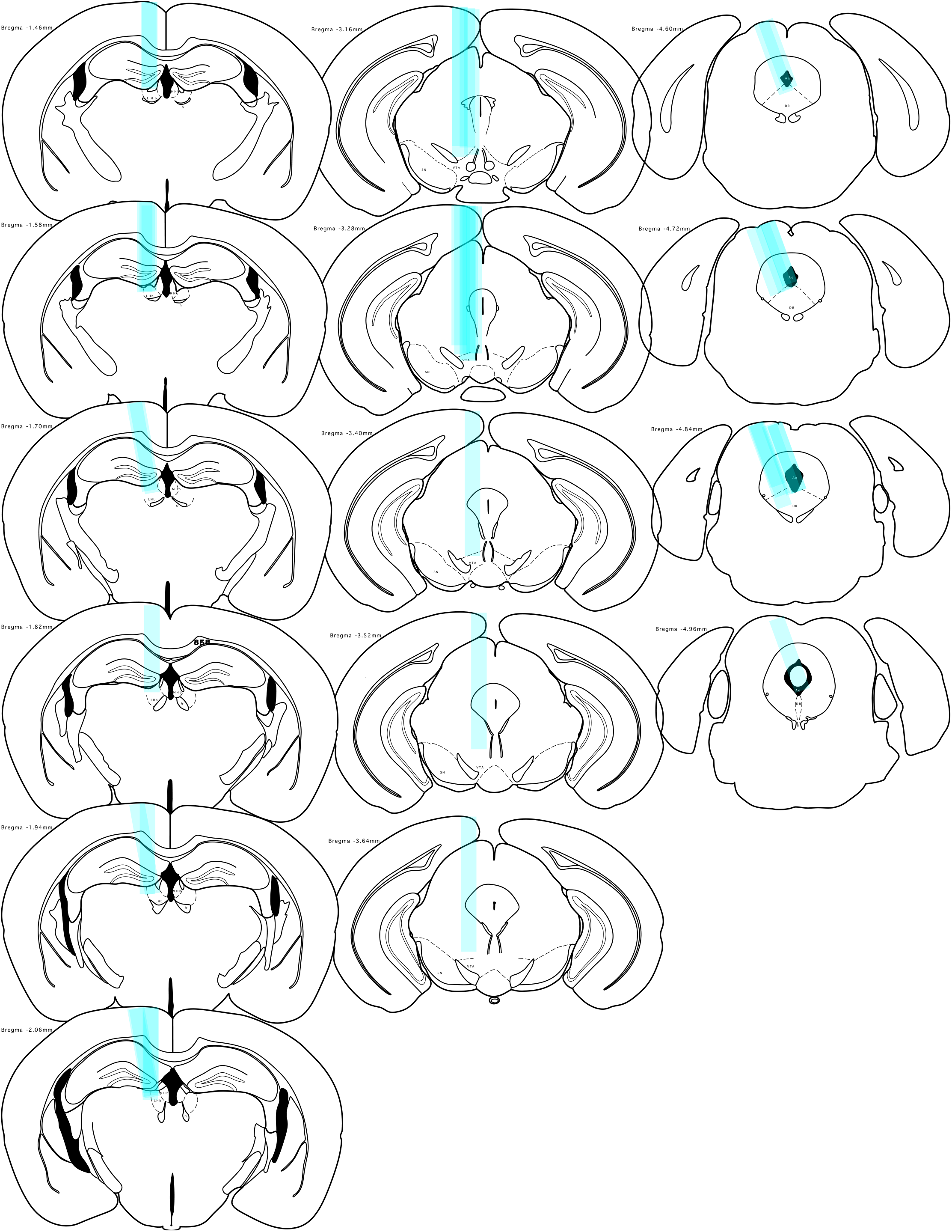
Cannulae placement in mice expressing GCaMP6s in the LHb (**left**), the VTA (**middle**), and the DRN (**right**)

**Figure 1–Figure supplement 4.**
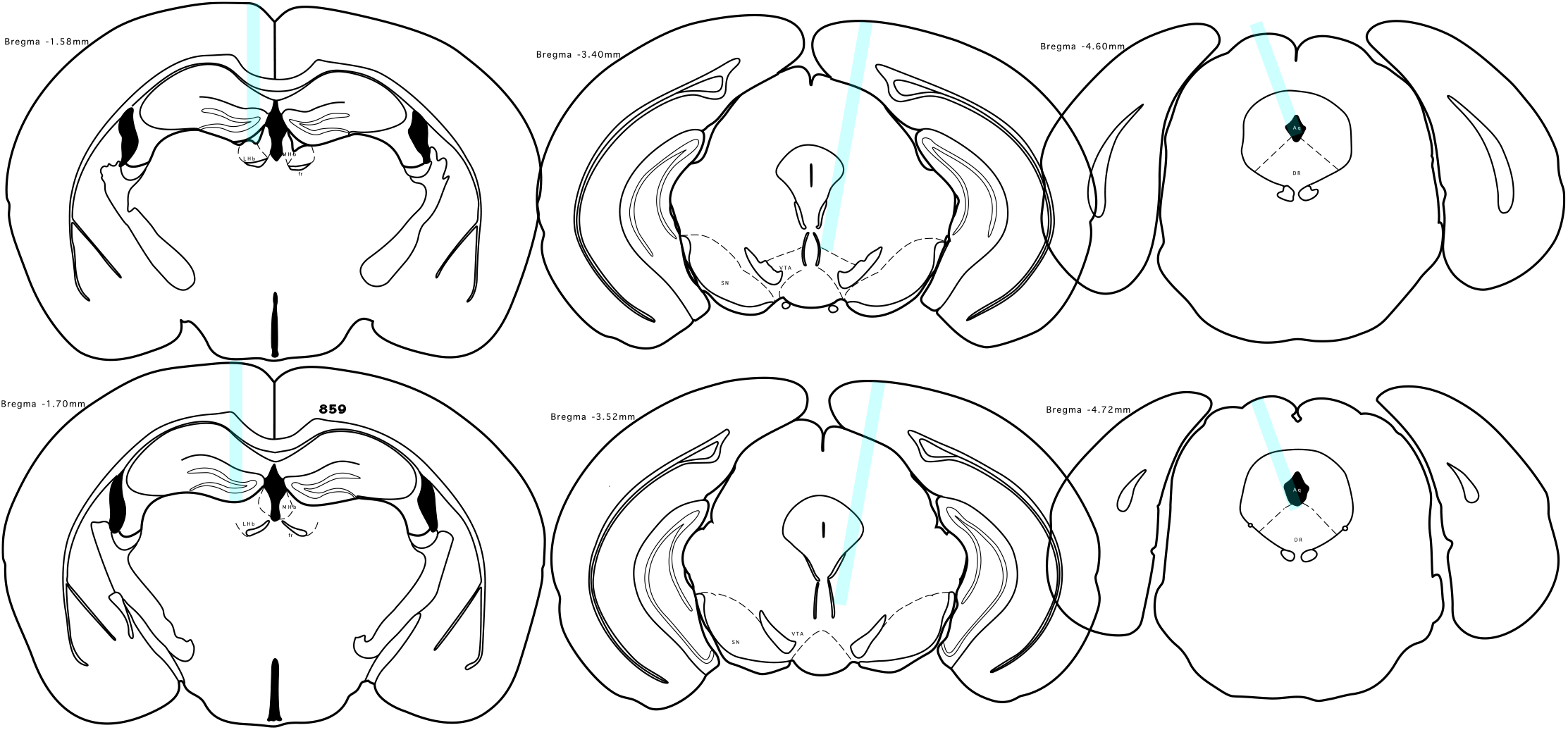
Cannulae placement in the mice expressing eYFP in the LHb (**left**), the VTA (**middle**), and the DRN (**right**)

**Figure 1–Figure supplement 5.**
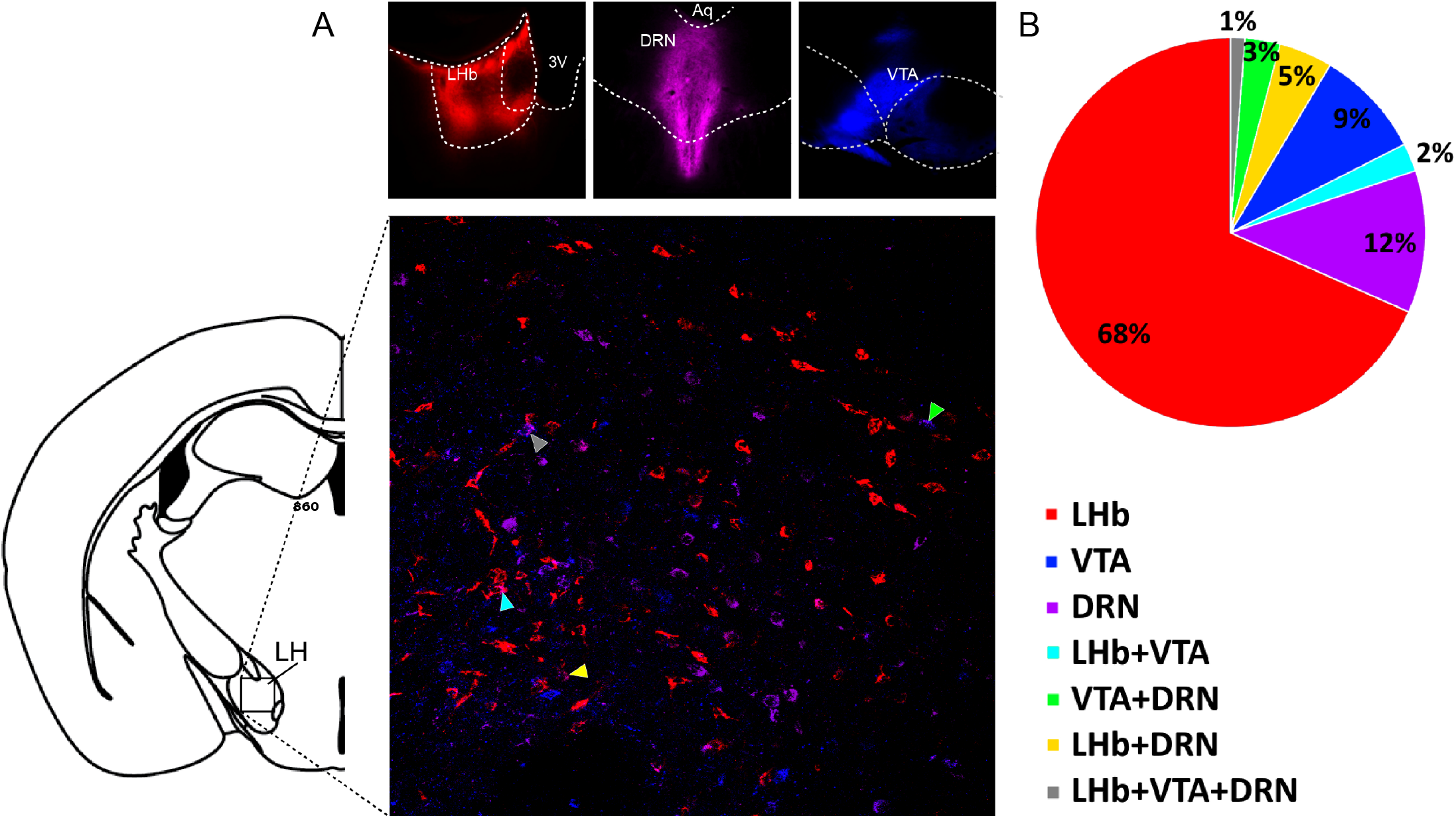
The LH neural populations projecting to the DRN, VTA, and LHb are largely distinct populations. (**A**) Representative fluorescent images of the fluorescent markers CTx 488, CTx 594, and CTx 647 (CTx – Cholera toxin) injected in the DRN, the VTA, and the LHb (top panels), and representative confical image of LH neurons positive for markers (bottom panel). Images are pseudo colored. (**B**) Pie chart representing the relative fraction of labeled LH neurons projecting to either the DRN, VTA, or LHb, and their co-localization.

**Figure 1–Figure supplement 6.**
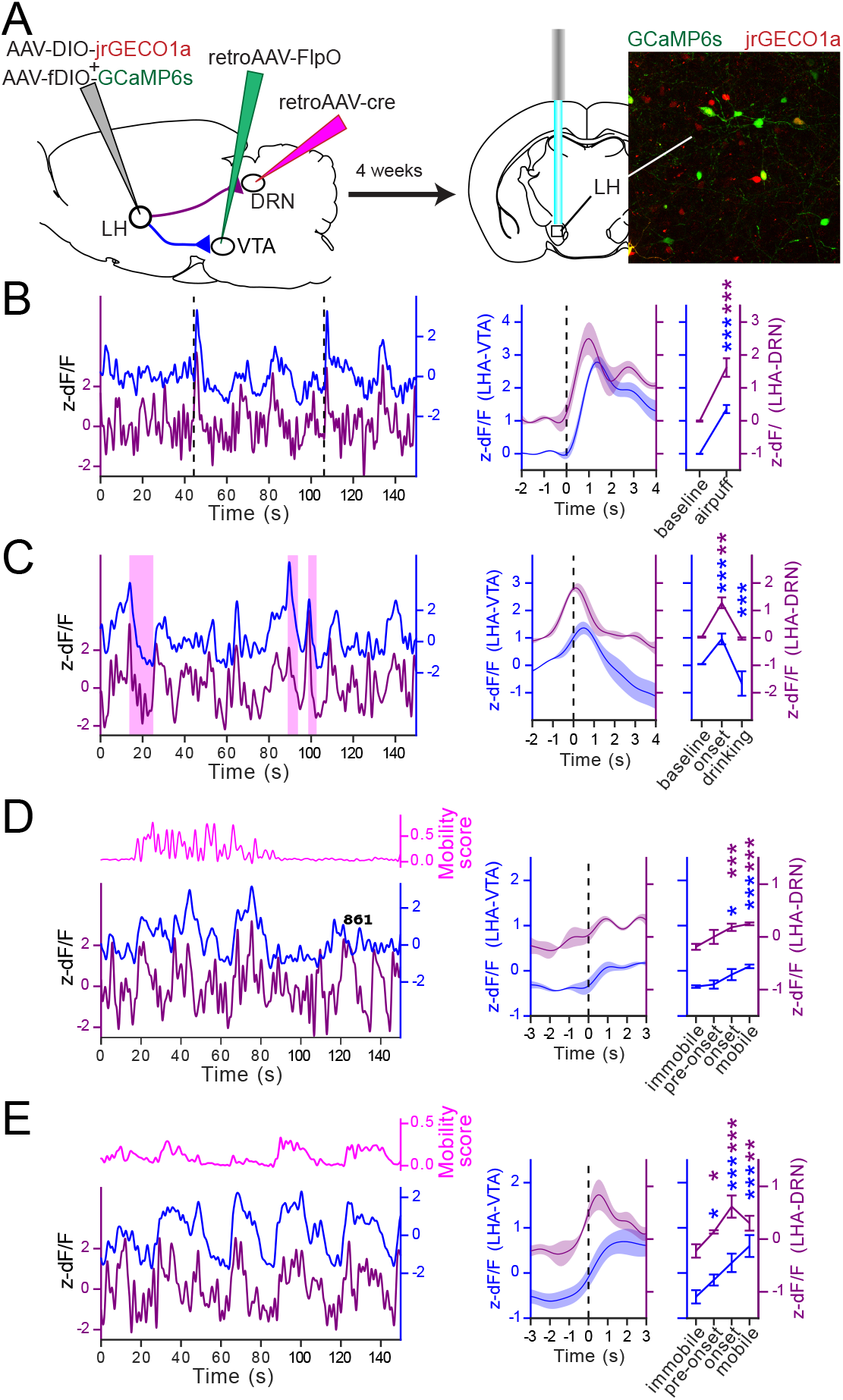
*Ca*^2+^ imaging from the LH neurons projecting to the DRN and the VTA. (**A**) Diagram of the experimental setup and representative fluorescent image of LH neuron expressing GCaMP6s (green) and jrGECO1a (red) projecting to the VTA and DRN, respectively. (**B**) Representative *Ca*^2+^ signal traces associated with the airpuffs (dashed vertical bars) simultaneously measured from the LH neurons expressing GCaMP6s (projecting to the VTA) and jrGECO1a (projecting to the DRN) (**left**). Peri-event plot of the average *Ca*^2+^ signals to all the airpuff events at the LH→DRN and LH→VTA LH neurons, and plot of the average responses before and after the airpuffs (**right**). Lines represent the means ± SEM. Same convention as for **B** for the sucrose consumption test (**C**), the open field test (**D**), and the tail suspension test (**E**). The sucrose consumption events are represented by the pink shaded boxes in **C**. The magenta lines are the mobility scores. Repeated measures two-way ANOVA within factors pathways (LH→DRN and LH→VTA) and time periods (different for each test) with post hoc Dunnett’s test. The *p* values were adjusted using the Bonferroni multiple testing correction method. \**p* < 0.05, \*\**p* < 0.01, \*\*\**p* < 0.001.

**Figure 1–Figure supplement 7.**
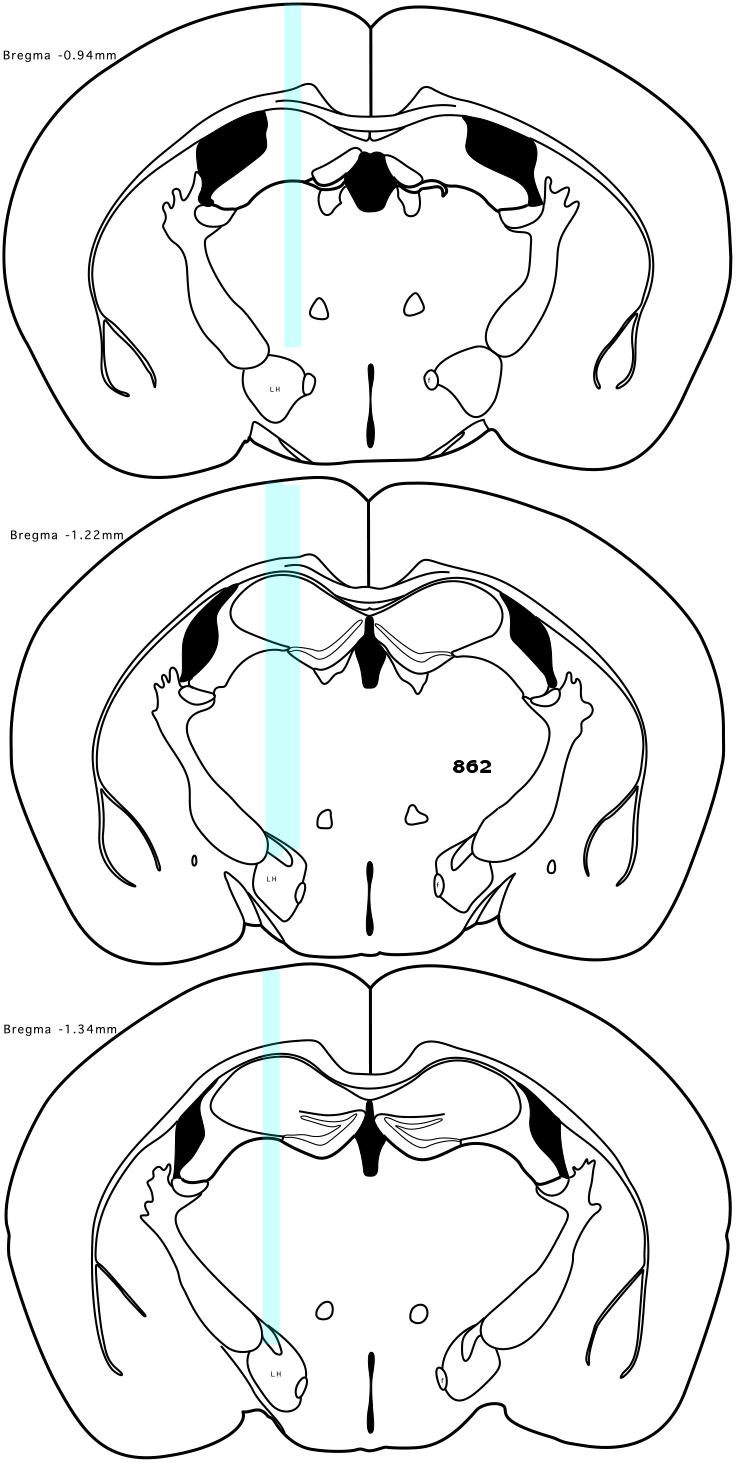
The cannulae placement in mice prepared using the intersectional viral strategy

**Figure 2–Figure supplement 1.**
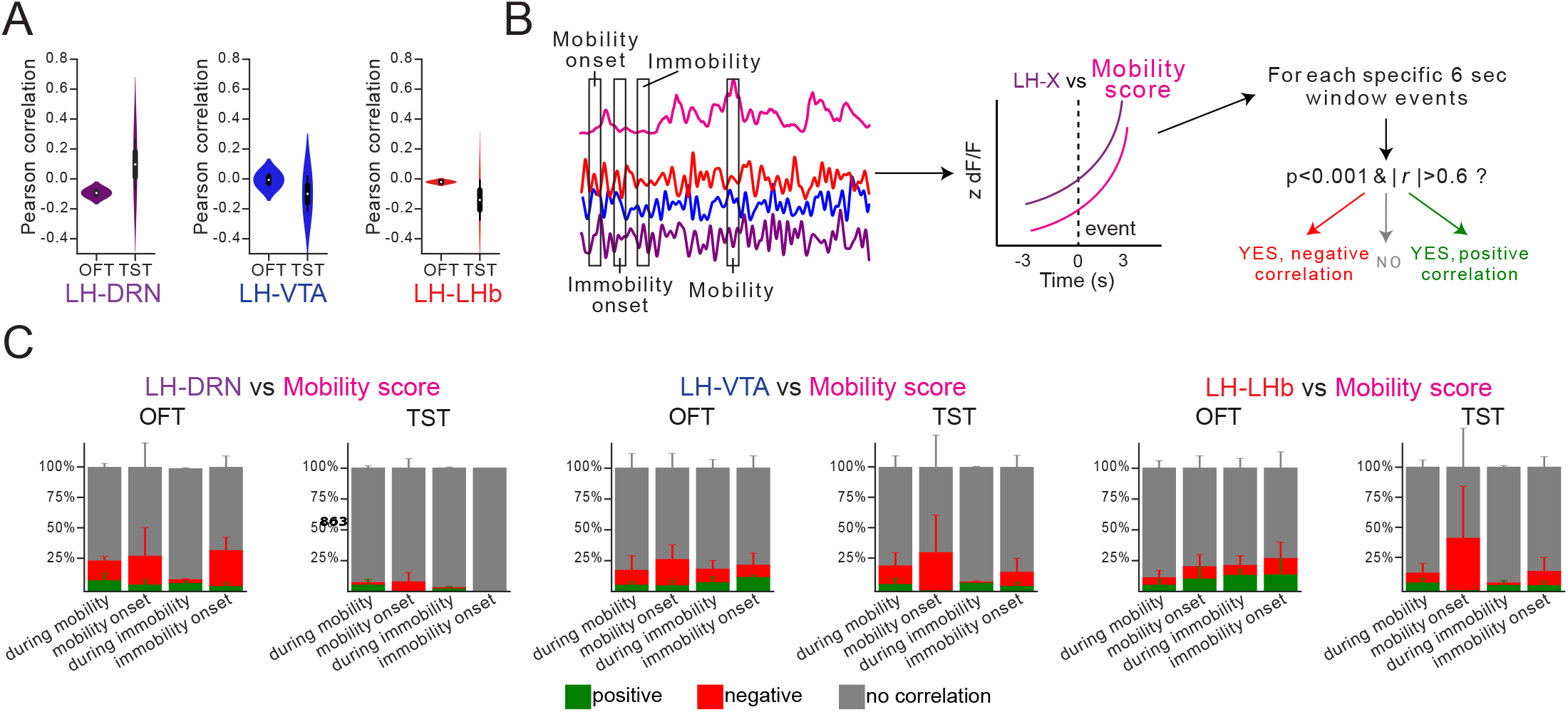
Correlation analysis between the signal at one of the LH neural output pathways and mobility score in mice expressing eYFP.. (**A**) The Pearson correlation between the *Ca*^2+^ signal measured at the LH→DRN, LH→VTA, and LH→LHb pathways and the mobility score during a complete session in the OFT or the TST. (**B**) Schematic of the event selection. Events at the onset of mobility and immobility and random events during mobility and immobility were chosen, and the Pearson correlation at 6-seconds peri-events between the *Ca*^2+^ signal and the mobility score was calculated. Correlations with p < 0.001 and r > 0.6 were considered as positive, p < 0.001 and r < 0.6 – negative, the others – not correlated. (**C**) Fraction of positive (green), negative (red), and uncorrelated events (gray) in the OFT and TST for the LH→DRN, LH→VTA, and LH→LHb pathways. Statistical analysis was performed along with the data from the mice expressing GCaMP6s.

**Figure 3–Figure supplement 1.**
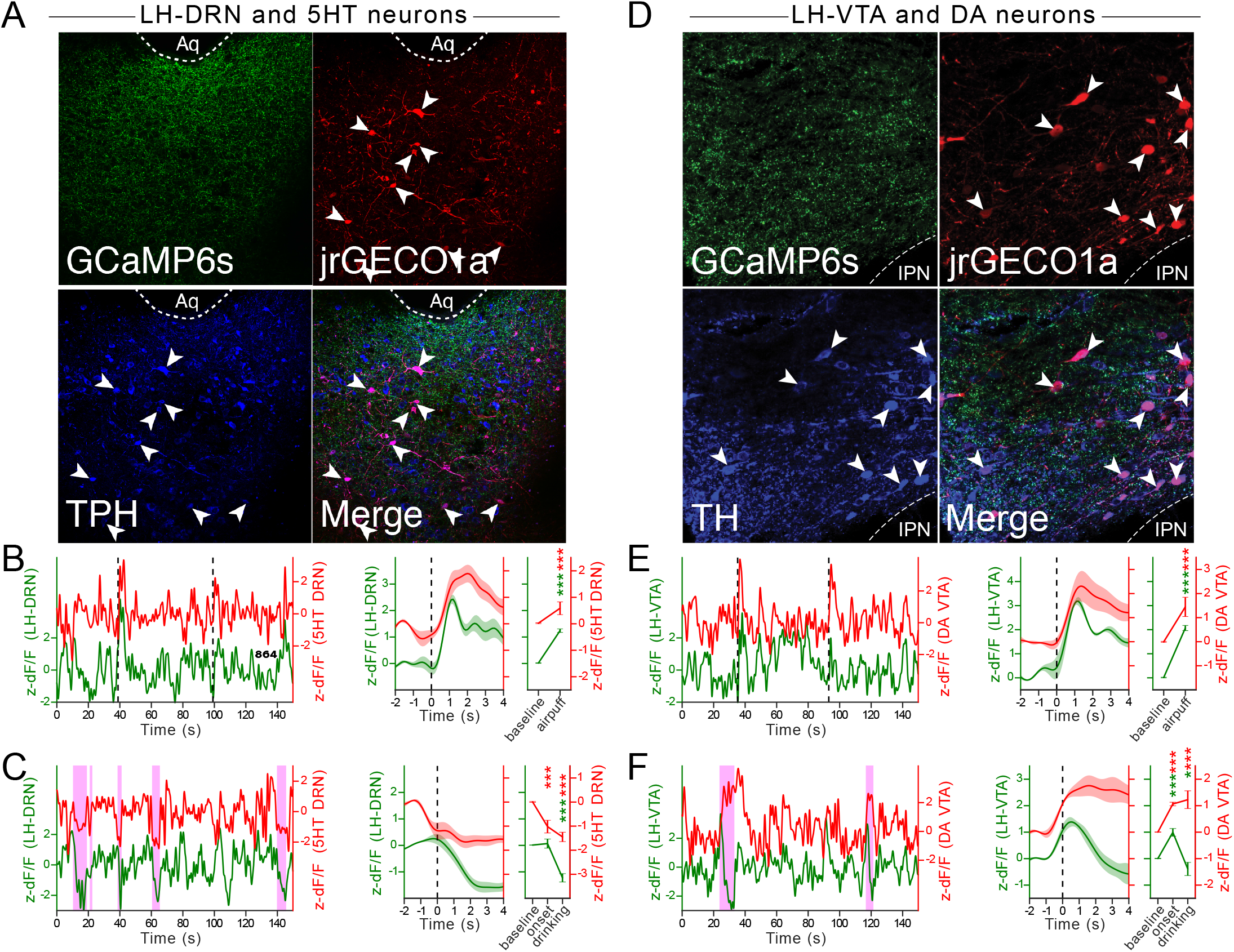
Simultaneous recordings at the DRN^5*HT*^ neurons and the LH→DRN pathway, and at the VTA^*DA*^ neurons and the LH→VTA pathway in the AP and the SCT. (**A, D**) Confocal images of the DRN neurons (**A**) and VTA neurons (**D**) expressing jrGECO1a and immunolabelled for the serotoninergic marker tryptophane hydroxylase (TPH, blue **A**) or the dopaminergic marker tyrosine hydroxylase (TH, blue **D**). Shown in green are the LH axon terminals in the DRN and VTA that are expressing GCaMP6s. Representative *Ca*^2+^ signal traces recorded from the DRN^5*HT*^ neurons and at the LH→DRN pathway (**B**) or from the VTA^*DA*^ neurons and at the LH→VTA pathway (**E**) in mice presented with airpuffs (**B, E**) (**left**). Peri-event plots of the average *Ca*^2+^ signal traces with all the onset of mobility and the plots for the AUC at baseline and after the airfpuffs (**right**). The lines represent means ± SEM. Same convention as for **B, E** for the SCT (**C, F**). Repeated measures two-way ANOVA within factors pathways (DRN^5*HT*^ and LH→DRN or VTA^*DA*^ and LH→VTA) and time periods (during immobility and mobility, at mobility pre-onset and onset) with post hoc Dunnett’s test. The *p* values were adjusted using the Bonferroni multiple testing correction method. \**p* < 0.05, \*\**p* < 0.01, \*\*\**p* < 0.001.

**Figure 3–Figure supplement 2.**
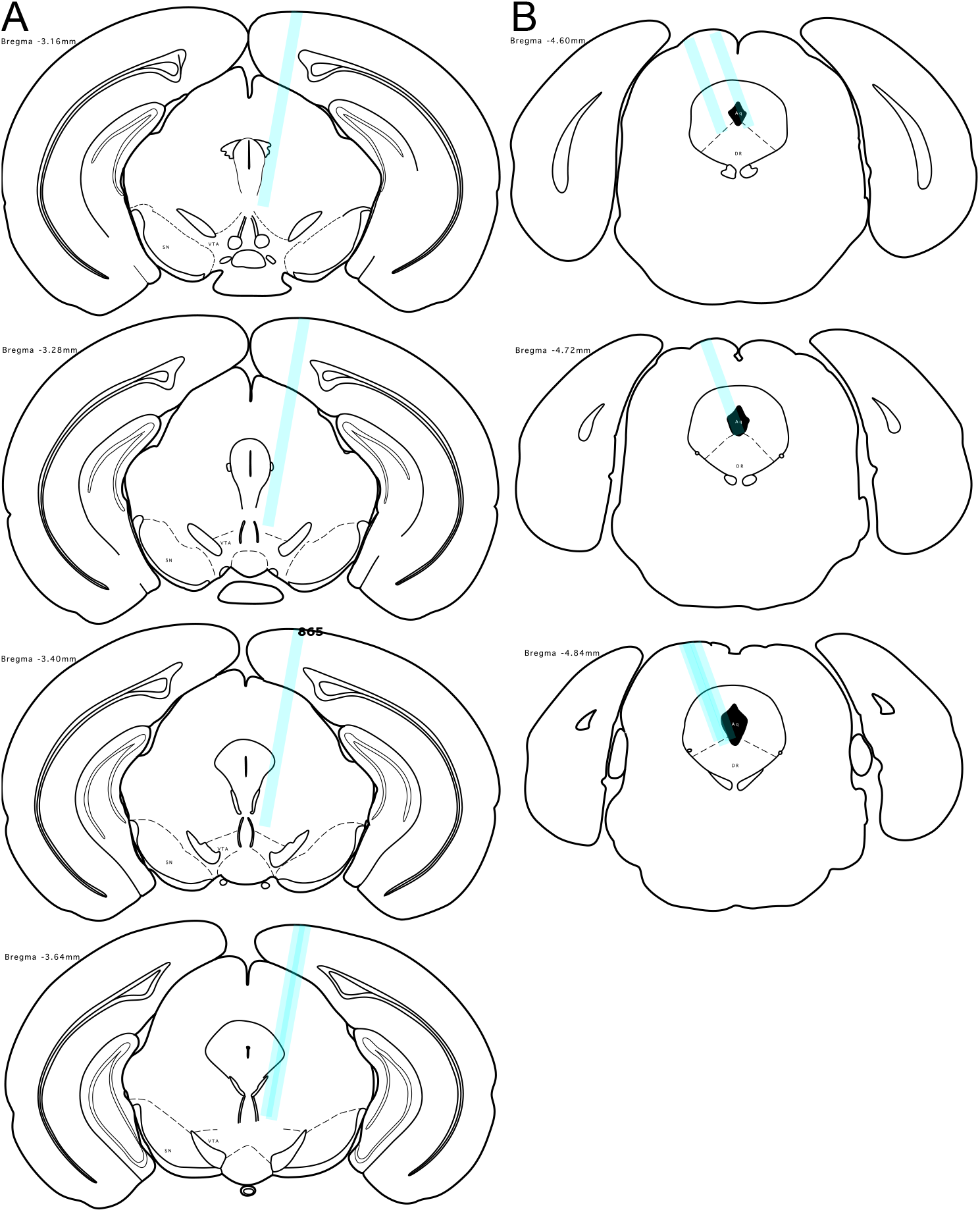
Cannulae placement in the ePet-cre (**A**) and DAT-ires-cre (**B**) mice

**Figure 4–Figure supplement 1.**
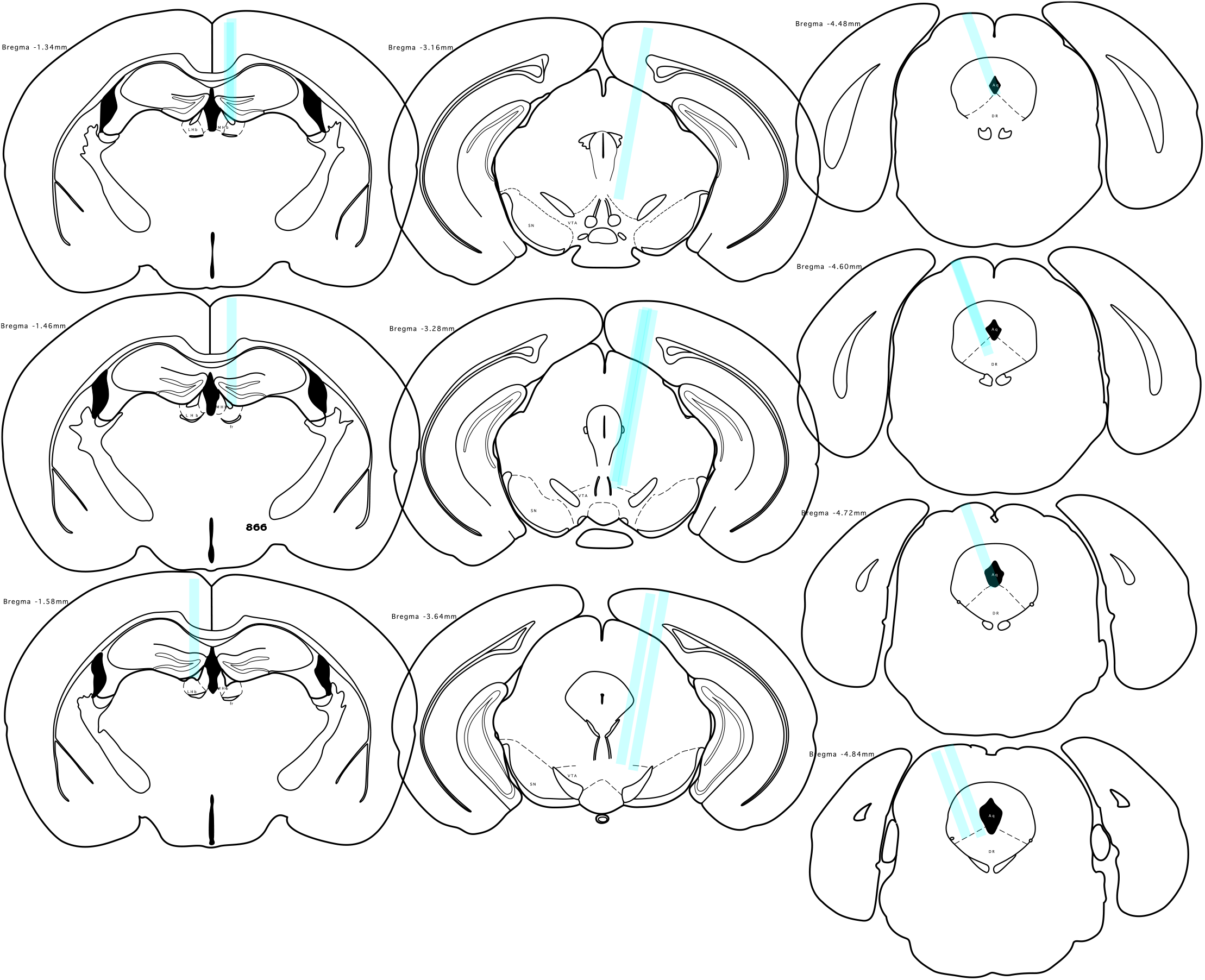
Cannulae placement in the LHb (**left**), the VTA (**middle**), and the DRN (**right**) of mice tested in the active avoidance task.

**Figure 6–Figure supplement 1.**
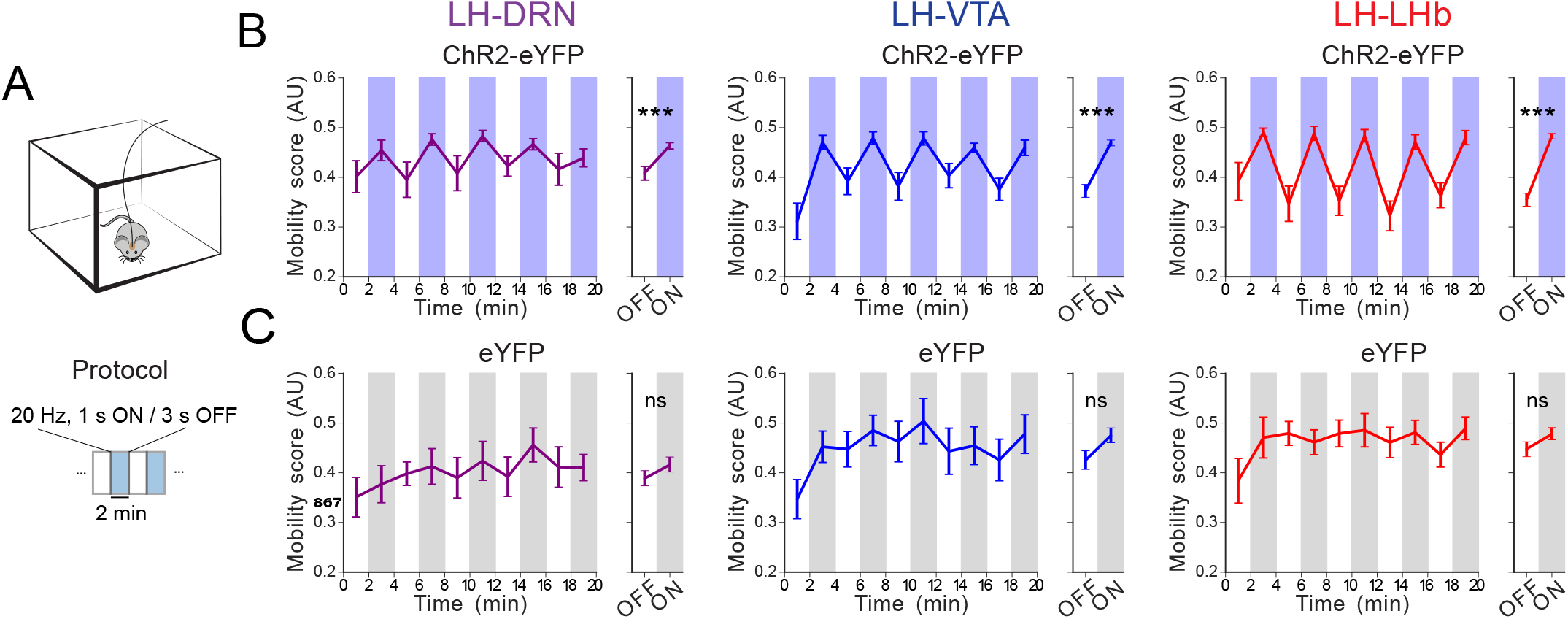
Optostimulation of the LH→DRN, LH→VTA, or LH→LHb pathways in the OFT. (**A**) Diagram of the experimental protocol for the OFT. The mobility of the mice was evaluated during a 20 min OFT session with alternating 2-min epochs without or with 1-s, 20 Hz trains every 4 sec. Mobility was automatically monitored with a video tracking system. (**B-C**) Plots of mean mobility (mean ± SEM) during periods of optogenetic stimulation (blue (**C**) or gray (**D**)) or no light (white) at the LH→DRN (**left**), LH→VTA (**middle**), or LHA→LHb (**right**) pathways in ChR2-eYFP- (**C**) and eYFP-expressing (**D**) mice. Four-way repeated measures ANOVA between factors of group (ChR2-eYFP- and eYFP-expressing mice), and within factors pathway (LH→DRN, LH→VTA, or LH→LHb), time period (five 4-minutes periods) and laser (on and off) with post hoc Tukey test. The *p* values were adjusted using the Bonferroni multiple testing correction method. \**p* < 0.05, \*\**p* < 0.01, \*\*\**p* < 0.001, *p* > 0.2 ns (not significant).

**Figure 6–Figure supplement 2.**
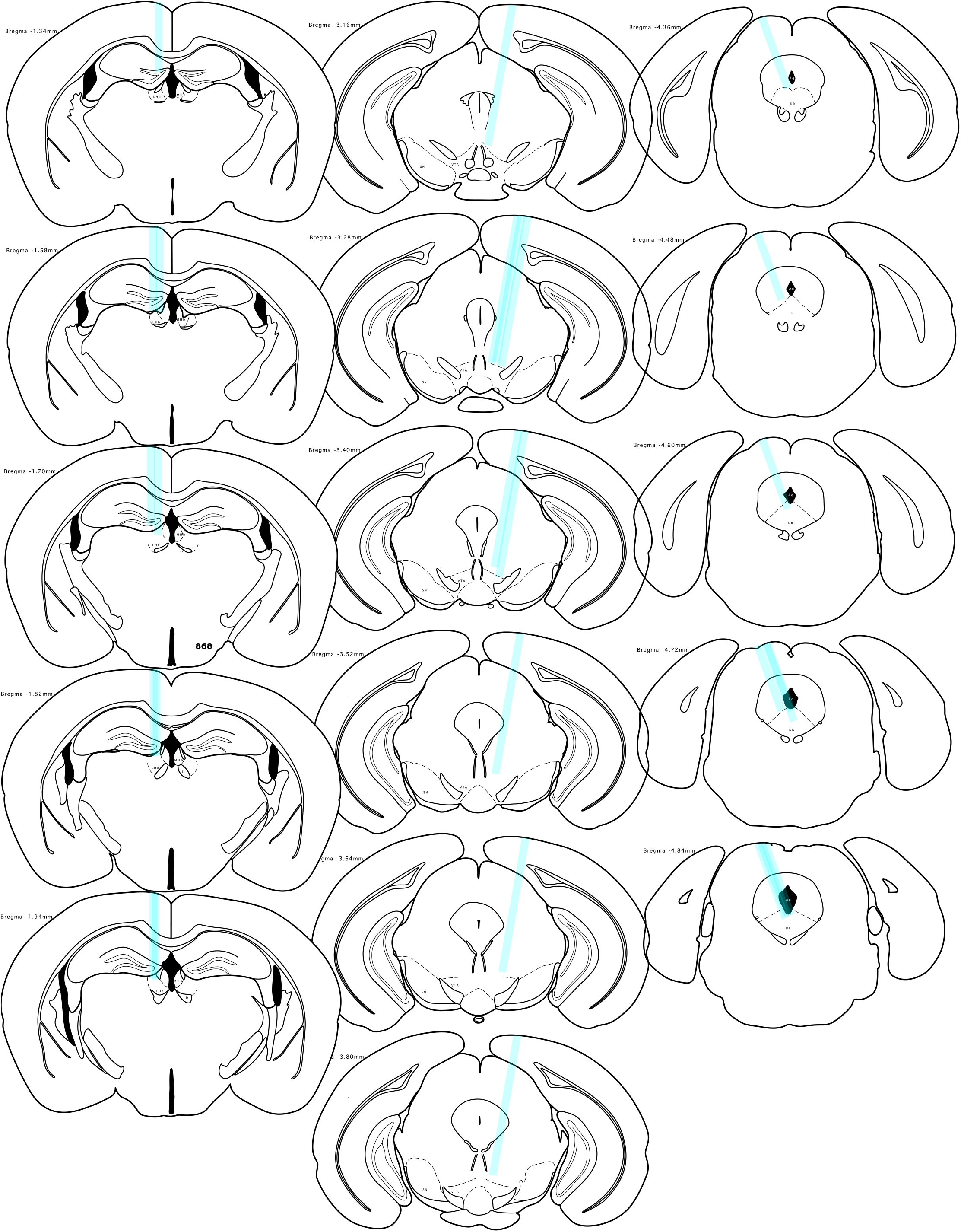
Cannulae placement in the LHb (**left**), the VTA (**middle**), and the DRN (**right**)in mice expressing ChR2-eYFP

**Figure 6–Figure supplement 3.**
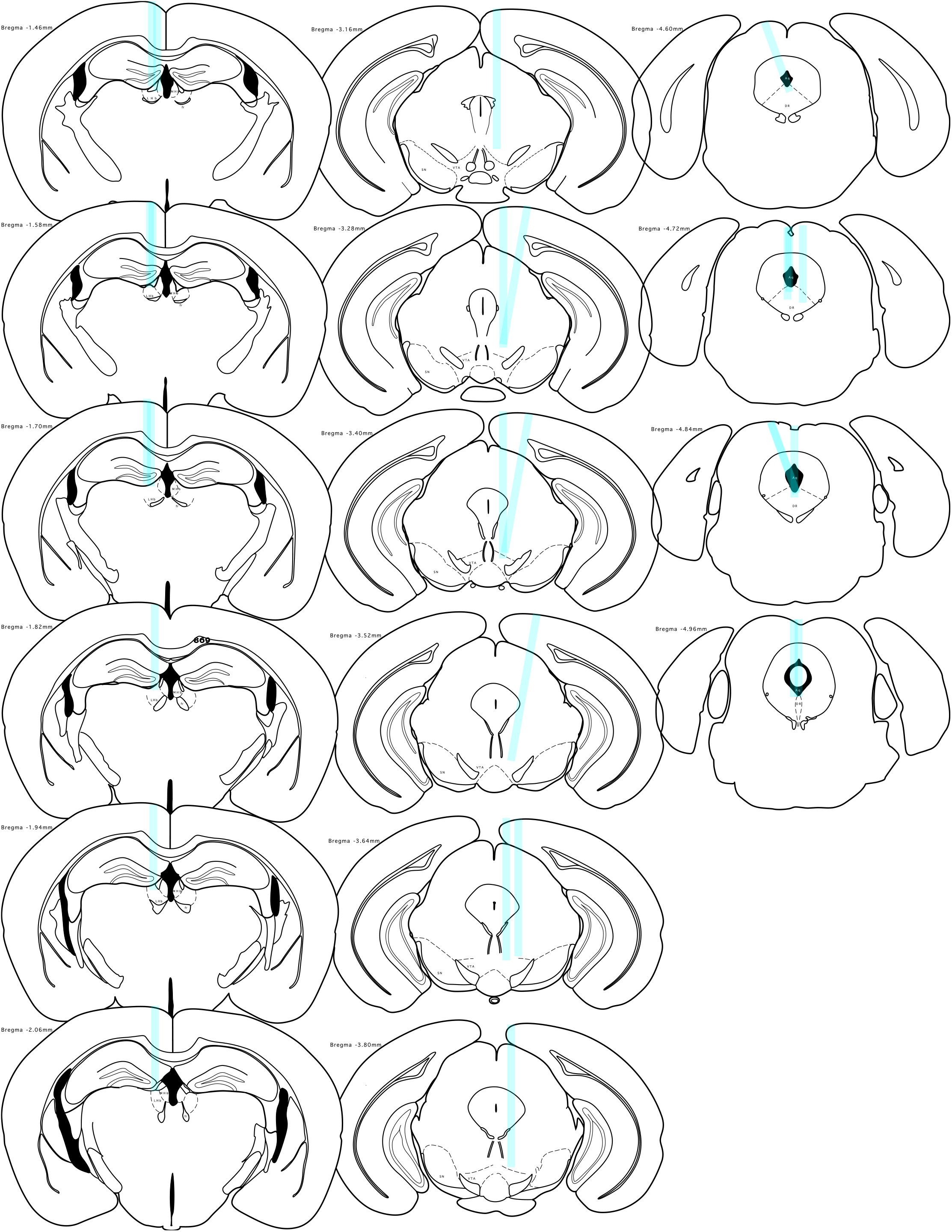
Cannulae placement in the LHb (**left**), the VTA (**middle**), and the DRN (**right**)in mice expressing eYFP

**Figure 7–Figure supplement 1.**
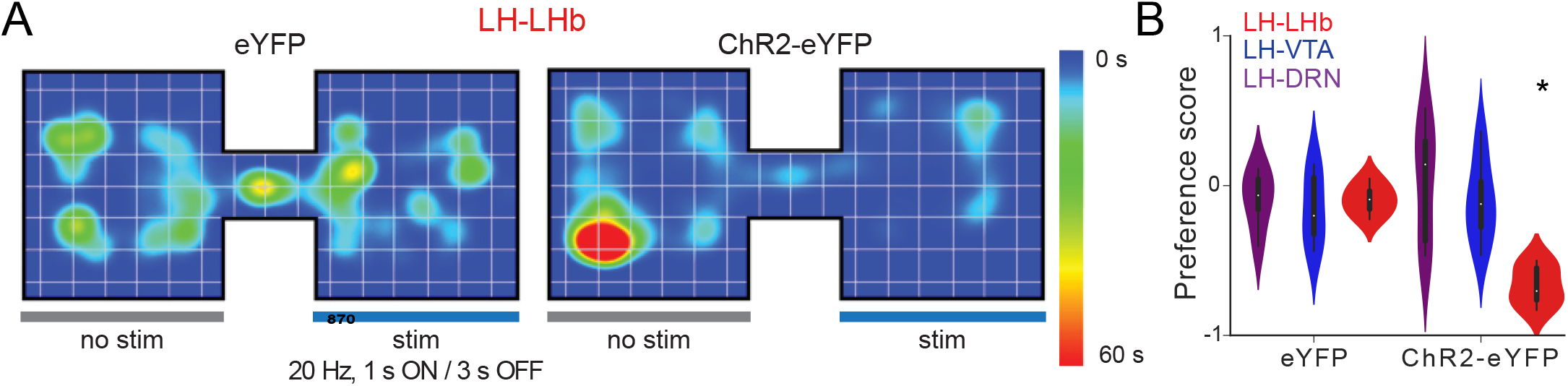
Optostimulation of the LH→DRN, LH→VTA, or LH→LHb pathways in the RTPP and the SCT. (**A**) The heatmaps of animal location in the RTPP during optostimulation of the LH→LHb pathway in mice expressing ChR2-eYFP (**left**) or eYFP (**right**). (**B**) The preference scores measured in the RTPP during optostimuation of the LH→DRN, LH→VTA, or LH→LHb pathways in mice expressing ChR2-eYFP or eYFP. Two-way ANOVA between factor of group (ChR2-eYFP- and eYFP-expressing mice) and within factor pathway (LH→DRN, LH→VTA, or LHA→LHb) with post hoc Tukey test. The *p* values were adjusted using the Bonferroni multiple testing correction method. #*p* < 0.1,\**p* < 0.05, \*\**p* < 0.01, \*\*\**p* < 0.001, ns (not significant) – *p* > 0.2.

